# A cytochrome P450 from juvenile mustard leaf beetles hydroxylates geraniol, a key step in iridoid biosynthesis

**DOI:** 10.1101/634485

**Authors:** Nanxia Fu, Zhi-ling Yang, Yannick Pauchet, Christian Paetz, Wolfgang Brandt, Wilhelm Boland, Antje Burse

## Abstract

Juveniles of the leaf beetle *Phaedon cochleariae* synthesize iridoid via the mevalonate pathway to repel predators. The normal terpenoid biosynthesis is integrated into the dedicated defensive pathway by the ω-hydroxylation of geraniol to 8-hydroxygeraniol. Here we identify and characterize the geraniol 8-hydroxylase as a P450 monooxygenase using integrated transcriptomic and proteomic analyses. In the fat body, 73 individual cytochrome P450s were identified. The double stranded RNA (dsRNA)-mediated knock down of *CYP6BH5* led to a significant reduction of 8-hydroxygeraniol-glucoside in the hemolymph and, later, of the chrysomelidial in the defensive secretion. Heterologously expressed CYP6BH5 converted geraniol to 8-hydroxygeraniol. In addition to geraniol, CYP6BH5 also catalyzes other monoterpenols, such as nerol and citronellol, into the corresponding α, ω-dihydroxy compounds.

**Highlights:**

- The geraniol 8-hydroxylase in *Phaedon cochleariae* was identified as a cytochrome P450 CYP6BH5.
- RNA interference emphasized the importance of CYP6BH5 in iridoid biosynthesis.
- *In vitro* enzyme assays showed that recombinant CYP6BH5 is a substrate promiscuous enzyme, converting the ω-hydroxylation of geraniol, nerol, citronellol but not linalool.
- Homology modeling suggested the -OH group of the substrate plays an important role in coordinating the substrates with the enzyme’s catalytic cavity.

## 1. Introduction

Iridoids comprise a large family of biologically active molecules that have so far been found in plants and insects. Structurally, they are known as *cis*-fused cyclopentan-[c]-pyrans with a hydroxyl (iridoid aglucones) or glucosyloxy group (iridoid glucosides) at C-1 position of the pyran ring. Besides their genuine defensive function against herbivores or insect predators, extensive studies have revealed that iridoids are also pharmaceutically valuable for the development of novel drugs and therapeutic strategies against diverse conditions including inflammation and cancers (Boros and Stermitz, 1991; Ghisalberti, 1998; Laurent et al., 2005; Rosa et al., 2008; Yamane et al., 2010). However, the often low yield of iridoids from natural resources has greatly hampered their therapeutic application (Miettinen et al., 2014). Maximizing the production of iridoids to an industrial scale using metabolic engineering offers a promising solution to their current scarcity (Alagna et al., 2016; Brown et al., 2015). To facilitate the optimization of the metabolic engineering, the identification of the genes and enzymes required for iridoid biosynthesis is essential.

Madagascar periwinkle, *Catharanthus roseus*, is a well-characterized iridoid-producing medicinal plant. Recently, all genes and enzymes involved in the iridoid biosynthetic pathway were completely elucidated in *C. roseus* (Krithika et al., 2015; Larsen et al., 2017; Miettinen et al., 2014). Although iridoids are typically encountered in the plant kingdom, these secondary metabolites have also been identified in insects (Cavill et al., 1984; Oldham et al., 1996; Smith et al., 1979). In fact, the name iridoid is a generic term derived from iridomyrmecin, iridolactone and iridodial, components of defensive secretions identified from species of the ant genus *Iridomyrmex* (Cavill et al., 1984). However, in insects, the iridoid pathway has been characterized in only a few species, and in *Iridomyrmex*, the pathway is even not yet fully resolved. The best-investigated insect species so far belong to the family of leaf beetles (Chrysomelidae). In particular, juveniles of Chrysomelina beetles, such as the mustard leaf beetle, *Phaedon cochleariae*, have evolved specialized pair-wise exocrine glands (composed of a reservoir with adhering glandular cells) to fend off predators; these glands, located on the beetle’s dorsal segment, are able to release defensive droplets containing iridoids. As shown in Figure 1, the iridoid *de novo* biosynthesis in *P. cochleariae* larvae starts with the formation of the early stage precursor 8-hydroxygeraniol glucoside (8-OH-Ger-Glc) in the fat body, which is transported via the hemolymph into the glandular reservoir (Burse et al., 2009, 2007; Frick et al., 2013; Strauss et al., 2013). In the glandular reservoir, 8-OH-Ger-Glc is subject to sequential hydrolysis, oxidation and cyclization to form the ultimate product, chrysomelidial (Bodemann et al., 2012; Rahfeld et al., 2015, 2014; Strauss et al., 2013). Although most of the glandular enzymes involved in the later steps of the pathway are well understood, to date only a few enzymes from the early biosynthetic steps have been functionally characterized (Burse et al., 2009; Frick et al., 2013; Kunert et al., 2013; Rahfeld et al., 2015; Strauss et al., 2013). Similar to other monoterpernoids, iridoid biosynthesis in *P. cochleariae* is initiated from geranyl pyrophosphate (GPP), a compound that is synthesized from the mevalonate pathway (Frick et al., 2013; Snyder and Qi, 2013). GPP is supposed to be further metabolized by a series of enzymes to generate the shuttling glucoside 8-OH-Ger-Glc.

**Figure 1.**
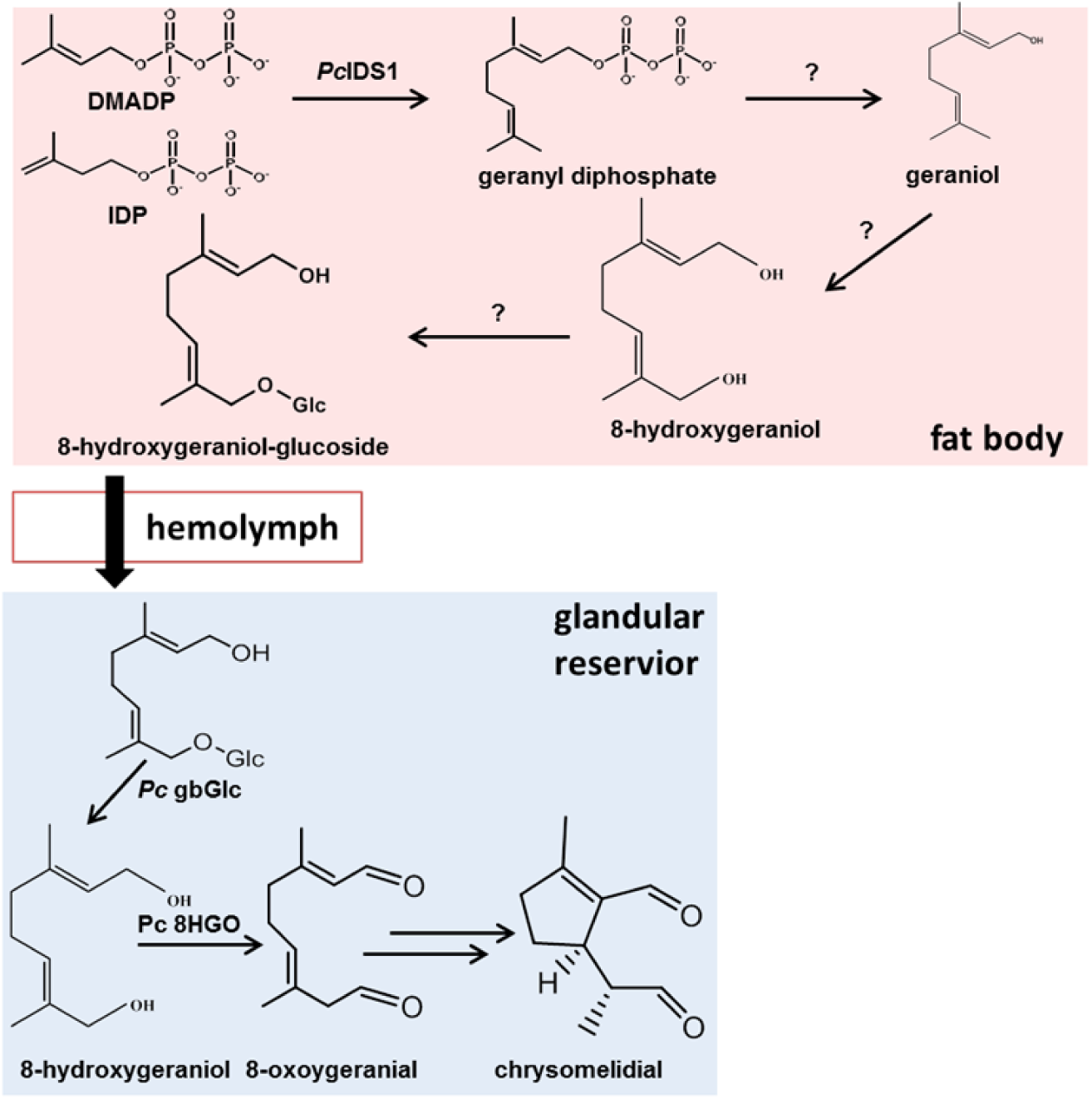
Iridoid biosynthetic pathway. Adopted and modified from Burse et al. (Burse et al., 2007).

Stable isotope labeling studies indicated the production of shuttling glucoside 8-OH-Ger-Glc in *P. cochleariae* was likely to proceed as follows (Oldham et al., 1996; Veith et al., 1994). First, the phosphate group is first removed from GPP by a presumable phosphatase, yielding geraniol (Burse et al., 2007). Afterwards, an oxidase converts geraniol into the diol 8-hydroxygeraniol (8-OH-geraniol) (Veith et al., 1994). In the step that follows, a glucosyltransferase catalyzes the addition of a sugar moiety to form the desired shuttling glucoside (Kunert et al., 2013). In *C. roseus*, the hydroxylation from geraniol to 8-OH-geraniol is catalyzed by a cytochrome P450 (Collu et al., 2001). It is hypothesized that the enzymatic system that produces 8-OH-geraniol in juvenile *P. cochleariae* has similar characteristics as in plants. However, the molecular characteristics of such an enzyme remain elusive.

Using an integrated approach coupling transcriptomic, proteomic and RNAi, herein, we identified a cytochrome P450, CYP6BH5, in juvenile *P. cochleariae*, which proved to be one of the missing enzymes in the early steps of iridoid biosynthesis. Substrate specificity analysis demonstrated that CYP6BH5 is a promiscuous enzyme, and the newly identified P450 can serve as an alternative catalyst for the production of iridoid via metabolic engineering.

## 2. Materials and methods

### 2.1. Leaf beetles

The mustard leaf beetle, *P. cochleariae*, was reared in the lab on Chinese cabbage *Brassica oleracea* convar. *capitata* var. *alba* (Gloria F1) under 16 hours light to 8 hours dark cycle conditions at 15±2 °C. *Chrysomela populi* (L.) were collected near Dornburg, Germany (+51°00’52.00”, +11°38’17.00”), on *Populus maximowiczii* x *Populus nigra*. The beetles were then lab-reared under 18°C ± 2°C in light (16 hours) and 13°C ± 2°C in darkness (8 hours) cycle conditions.

### 2.2. RNA extraction and cDNA synthesis

Different tissues (fat body, gut, gland and Malpighian tubules) of *P. cochleariae* larvae were dissected under the microscope and stored in 100 µl lysis-buffer (Life Technologies, Carlsbad, CA, USA) with the addition of 1 µl ExpressArt NucleoGuard (Amp Tec GmbH, Hamburg, Germany) at −80 °C until needed. Total RNA from stored tissues was isolated with RNAqueous®-Micro Kit (Life Technologies, Carlsbad, CA, USA) according to the manufacturer’s instructions. The RNA from the larvae of *P. cochleariae* and *C. populi* was isolated with the RNAqueous kit (Life Technologies, Carlsbad, CA, USA). All RNA samples were stored at −80 °C after extraction. For RNA samples used for quantitative real-time PCR (qPCR), the genomic DNA was digested by DNase I (Thermo Fisher Scientific, Langenselbold, Germany) prior to cDNA synthesis.

For cDNA synthesis, a total of 400 ng RNA was used together with 1 µl SuperScript III Reverse Transcriptase (Life Technologies, Carlsbad, CA, USA) and 0.5 μg/μl Oligo(dT)_12-18_ Primers (Life Technologies, Carlsbad, CA, USA) per 20 μl reaction. The cDNA template from the *P. cochleariae* fat body for 3’ rapid amplification of cDNA ends (3’ RACE) was synthesized according to the manual from SMARTer RACE 5’/3’ Kit (Takara Bio, Inc. Mountain View, CA, USA). All cDNA templates were stored at −20 °C after synthesis.

### 2.3. Membrane proteomics and annotation of the cytochrome P450s in the fat body

To extract membrane protein, 100-200 mg fat body was used and separated via ultra-centrifugation at 100,000g. The membrane fractions were dissolved in 5x SDS sample buffer (50 mM Tris-HCl pH 6.8, 2% SDS, 10% glycerol, 1% *β*-mercaptoethanol, 12.5 mM EDTA, 0.02 % bromophenol blue) and stored at −20 °C until needed. Proteins of the membrane fractions were separated by SDS PAGE and stained afterwards with Coomassie blue. Proteomic analysis was performed by liquid chromatography-mass spectrometry (LC-MS^E^) according to Rahfeld et al. (2015). The proteomic data were processed with our in-house *P. cochleriae* transcriptome database (Stock et al., 2013). Afterwards, Pfam and Blast2Go analysis were used to identify all fat body cytochrome P450s.

### 2.4. Analysis of the expression pattern in selected tissues

Reverse transcription-qPCR (RT-qPCR) was performed to analyze the expression pattern of selected P450 candidates in different tissues (fat body, gut, Malpighian tubules and glands) with Bio-Rad CFX96 real time PCR detection system (Bio-Rad Laboratories, Munich, Germany). Each RT-qPCR reaction (20 μl final volume) contained 10 μl Brilliant III SYBR Green qPCR Master Mix (Agilent Technologies, Waldbronn, Germany), 0.6 μl of cDNA, and 0.2 μl each of forward and reverse gene-specific primers (Table S3). The initial incubation took place at 95 °C for 3 min, followed by 40 cycles at 95 °C for 10 s, 60 °C for 20 s, and at 72 °C for 30s. Three technical replicates were applied to three biological replicates. Technical replicates with a Cq difference >0.5 were excluded. Elongation factors 1a (*Pc* EF1a) and the eukaryotic translation initiation factor 4a (*Pc* eIF4a) were chosen as reference genes for normalization. Primers were designed by primer3plus (http://www.bioinformatics.nl/cgibin/primer3plus/primer3plus.cgi); all primer sequences can be found in Table S3. All assays were performed according to MIQE-guidelines (Bustin et al., 2009).

### 2.5. Silencing of candidate genes by RNA interference (RNAi)

The assembled sequences of selected candidates were analyzed by siFi 21 (https://sourceforge.net/projects/sifi21/) for off-target prediction using a threshold value of at least 21 continuous nucleotides. The unique fragments (>400 bp) were used to design dsRNA primers (Table S3) and all dsRNA fragments (including the ds *eGFP* control) were synthesized using the MEGAscript RNAi Kit (Thermo Fisher Scientific, Langenselbold, Germany). After purification, the dsRNA was diluted in 0.9% NaCl solution and adjusted to a concentration of 2 μg/µl. Second instar *P. cochleariae* larvae were anesthetized on ice prior to injection. For each larva, 100 ng dsRNA was injected into the hemolymph under the microscope with a glass capillary. Larvae injected with only dsRNA targeting *eGFP* were used as a control group. After injection, the larvae were transferred into insect rearing cups and kept under normal rearing conditions.

### 2.6. Collection of hemolymph and the glandular secretion

Larval glandular secretion and hemolymph were collected by glass capillaries (i.d.: 0.28 mm, o.d.: 0.78 mm, length: 100 mm; Hirschmann, Eberstadt, Germany) and then transferred to a 200 μl Eppendorf tube. The weight of the glandular secretion and hemolymph were obtained by weighing empty and filled tubes with hemolymph or secretion. Samples were stored at −20 °C until needed.

### 2.7. Amplification and sub-cloning of the *PcC7758* open reading frame (ORF)

To obtain the full-length ORF, RACE-PCR was conducted with combination of fat body 3’RACE ready cDNA and RACE primers (Table S3) according to the manufacturer’s protocol (Takara Bio, Mountain View, CA, USA). The ORF was obtained by assembling the RACE fragment and the corresponding fragment obtained from the transcriptome. Afterwards, the ORF sequence was re-amplified with Phusion High-Fidelity DNA Polymerase (Life Technologies, Carlsbad, CA, USA) and the corresponding primers (Table S3). The complete ORF sequence was verified by sequencing.

To produce the recombinant protein, *CYP6BH5* was sub-cloned into pcDNA™3.1D/V5-His-TOPO and pFastBac Dual expression vectors, respectively, according to the protocol (Thermo Fisher Scientific, Langenselbold, Germany). To facilitate detection of the recombinant proteins, constructs without a stop codon were sub-cloned in parallel into corresponding expression vectors. A *C. populi* cytochrome P450 reductase (*CPR*) was cloned and subjected to operations similar to those mentioned above. All primers for sub-cloning can be found in Table S3.

### 2.8. Expression of the recombinant proteins and microsome isolation

To produce recombinant proteins in HEK cells, the recombinant pcDNA™3.1D/V5-His-TOPO plasmids of *CYP6BH5* and *CPR* were co-transfected in a ratio of 4:1 into HEK cells by electroporation. HEK cells were maintained at 37 °C with 10% CO_2_. After 48 h, cells were harvested and washed with phosphate-buffered saline (PBS) buffer, and the microsomal fraction was prepared as described by Joussen et al. (2012) and stored at −80 °C. To produce recombinant proteins with Sf9 cells, the bacmids were transfected into the Sf9 insect cells using a Bac-to-Bac baculovirus expression system according to the manual (Thermo Fisher Scientific, Langenselbold, Germany). The titer of the recombinant virus was determined following manufacturer’s instructions. Sf9 cells were co-infected with recombinant baculoviruses expressing CYP6BH5 and CPR with a multiplicity of infection (MOI) of 1 and 0.1, respectively. Sf9 cells were maintained at 27 °C with Sf-900 II SFM medium (Life Technologies, Carlsbad, CA, USA), supplemented with 2.5 μg/ml hemin and 0.3% (vol/vol) fetal bovine serum (Atlas Biologicals, Fort Collins,CO, USA). After 72 h, cells were harvested for isolation of the microsomal fraction. The microsomal fraction was aliquoted and stored at −80 °C after protein quantification with the Quick Start Bradford Protein Assay Kit (Bio-Rad Laboratories, Munich, Germany).

### 2.9. SDS PAGE and western blot

Microsomal fractions containing CYP6BH5 and CPR, fused with a V5 epitope tag and a His-tag were denatured by incubation at 70 °C for 5 minutes and followed by SDS PAGE separation. Afterwards, membrane proteins were blotted onto a polyvinylidene difluoride (PVDF) membrane (Trans-Blot^®^ Turbo™ Mini PVDF Transfer Pack; Bio-Rad) with a Bio-Rad blotting system. The membrane was blocked first in blocking buffer (5% (wt/vol) non-fat dry milk in Tris buffered saline with Tween-20 (TBST) buffer) for 1 h and then incubated overnight with Anti-V5-HRP antibody (1:10,000; Thermo Fisher Scientific, Langenselbold, Germany) in another blocking buffer (0.25 % (wt/vol) non-fat dry milk in TBST buffer) at 4 °C. The membrane was washed three times with TBST buffer prior to incubation (1 min) with enhanced chemiluminescence (ECL) solution (Thermo Fisher Scientific, Langenselbold, Germany). The Amersham Hyperfilm ECL X-ray film (GE Healthcare, Boston, USA) was exposed to the PVDF membrane prior to developing the film.

### 2.10. Enzyme assays

The microsomal fraction with the recombinant proteins lacking the V5 epitope and His-tag was used for the enzyme assay. The assay was performed in 50 µl volume containing 25 μg microsomal proteins in 20 mM potassium phosphate buffer (pH 7.5), 200 µM substrate and 1 mM NADPH. The same assay without NADPH served as the negative control. The empty control, in contrast, contained 25 μg microsomal proteins obtained from cells that were infected with only the recombinant CPR vector was used. After incubation at 30 °C for 15 min, the reaction was quenched and extracted by the addition of an equal volume of ethyl acetate. Samples were silylated by N-methyl-N-(trimethylsilyl) trifluoroacetamide (MSTFA) prior to gas chromatography–mass spectrometry (GC-MS) analysis.

To generate products for nuclear magnetic resonance (NMR) analysis, the standard enzyme assay was scaled up to a volume of 500 µl containing 1 mM of substrate (nerol or citronellol). After a first round of incubation with 70 µg microsomes for 1 hour at 30 °C, a second aliquot of 70 µg microsomes was added and incubated for another 1 hour, and those steps were repeated 5 times. The reaction was then stopped by adding 200 µl of 1 M HCl, vortexing, and cooling on ice. Two up-scaled assays for each substrate were pooled. To extract the products, equal volumes of ethyl acetate were added and the supernatants were collected. The solvent of the samples was subsequently removed by a stream of nitrogen gas.

### 2.11. GC-MS, high performance liquid chromatography-mass spectrometry (HPLC-MS) and NMR analysis

To detect chrysomelidial, the stored larval secretion was dissolved in 25 μl ethyl acetate spiked with 2 μg/μl methyl benzoate, and the supernatant was transferred to GC vials. One microliter was subjected to GC-MS analysis [ThermoQuest ISQ mass spectrometer EI LI system (quadrupole) equipped with Phenomenex ZB-5-W/Guard Column, 25 m (10 m Guard Pre-column) × 0.25 mm, film thickness of 0.25 μm]. The program setting was the same as previously reported (Rahfeld et al., 2014). The peak areas were calculated by Thermo Xcalibur Quan Brower that is implemented in the Xcalibur software (Thermo Fisher Scientific, Langenselbold, Germany). The relative amount of chrysomelidial per µg secretion was calculated against the amount of methyl benzoate spiked in the solvent.

To detect the 8-OH-Ger-Glc, the stored larval hemolymph was extracted with 50 µl methanol containing 0.025 mM 8-hydroxygeraniol-8-thio-*β*-D-glucoside (Ger-8-S-Glc). The supernatant was collected and subjected to HPLC–MS analysis. Condition for the measurement and identification of 8-OH-Ger-Glc and Ger-8-S-Glc followed the method optimized by Kunert et al. (2008). The peak areas were calculated as mentioned earlier. The relative amount of 8-OH-Ger-Glc per µg hemolymph was calculated against the internal standard Ger-8-S-Glc.

For the detection of 8-OH-geraniol by GC-MS in extracts and enzyme assays, the hydroxyl groups of the analytes were first silylated with MSTFA at 70 °C for 30 minutes. Then, the sample was dried under nitrogen and re-suspended in ethyl acetate. One microliter was used for GC-MS analysis under programmed conditions: 50 °C (2 min), 10 °C/min to 280 °C, 30 °C/min to 310 °C (1 min). The inlet temperature was 230 °C and the transfer line temperature was 280 °C. 8-OH-geraniol was identified by comparing the retention time and mass spectra with an authentic standard. The structure of 8-hydroxynerol and 8-hydroxycitronellol was determined by NMR. ^1^H, ^1^H-^1^H COSY, ^1^H-^13^C HSQC, ^1^H-^13^C HMBC and ^1^H-^1^H ROESY were acquired on a 700 MHz Avance III HD spectrometer equipped with a 1.7 mm cryoprobe (Bruker Biospin, Germany). Data acquisition and processing were accomplished using TopSpin ver. 3.2 (Bruker Biospin, Rheinstetten, Germany). Samples were measured in CDCl_3_ at 293 K.

### 2.12. Homology modeling

The 3D-protein structure homology modeling of CYP6BH5 was performed with YASARA Version 17.12.24 (Krieger et al., 2009; Krieger and Vriend, 2014). After searching for templates in the protein database, ten appropriate X-ray templates were found (PDB-codes: 5VCC, 5VEU, 6C93, 5XA3, 6C94, 1ZOA, 3MDM, 5B2X, 2FDV) (Berman et al., 2000). A homology-modeling template was generated for each of these 10 X-ray templates based on alternative sequence alignment to 78 models. The quality of all homology models was evaluated using PROCHECK and ProSA II (Laskowski et al., 1993; Sippl, 1993, 1990). The model based on the X-ray structure of 5VCC, scored −1.613, and was the best, with a sequence identity of 34.2% and a sequence similarity of 55.0% (BLOSUM62 score is > 0). This model was subsequently refined with the md-refinement tool of YASARA (20 simulated annealing runs for 500 ps). A model with excellent quality resulted, based on the Ramachandran plot: 91.5% residues were found in the most favored region, and two outliers were found only in the loop regions; all the ProSA energy graphs are in the negative range, and the calculated z-scores are in the range of natively folded proteins.

100 docking positions of geraniol were generated by docking geraniol at the active site close to heme center using MOE 2018.01 (https://www.chemcomp.com/). Out of several alternative docking positions, the fourth one was selected because of the proximity between the C_10_-methyl group of geraniol and the peroxide bound to heme. To confirm the stability of the arrangement, a second md-refinement run was performed.

## 3. Results

### 3.1. Presence of 8-OH-geraniol in the fat body of *P. cochleariae*

The aglycone 8-OH-geraniol is the first metabolite that links classical terpenoid biosynthesis to dedicated defense metabolism. Although the presence of 8-OH-Ger-Glc in fat body extracts from *P. cochleariae* has been reported by Burse et al. (2007), the presence of the corresponding aglycone in the fat body has not been validated. Therefore, the presence of 8-OH-geraniol in this tissue was checked first. As shown in Figure 2, the production of 8-OH-geraniol in the fat body was confirmed using authentic standards (mass spectra, see Fig. S1). Moreover, its glucoside could also be detected in the fat body with HPLC-MS (Fig. S2). This observations support the hypothesis that the early biosynthetic steps of the iridoid biosynthesis pathway are localized in the fat body (Burse et al., 2007), which therefore should contain corresponding enzymes including the geraniol oxidase and the glycosyltransferase.

**Figure 2.**
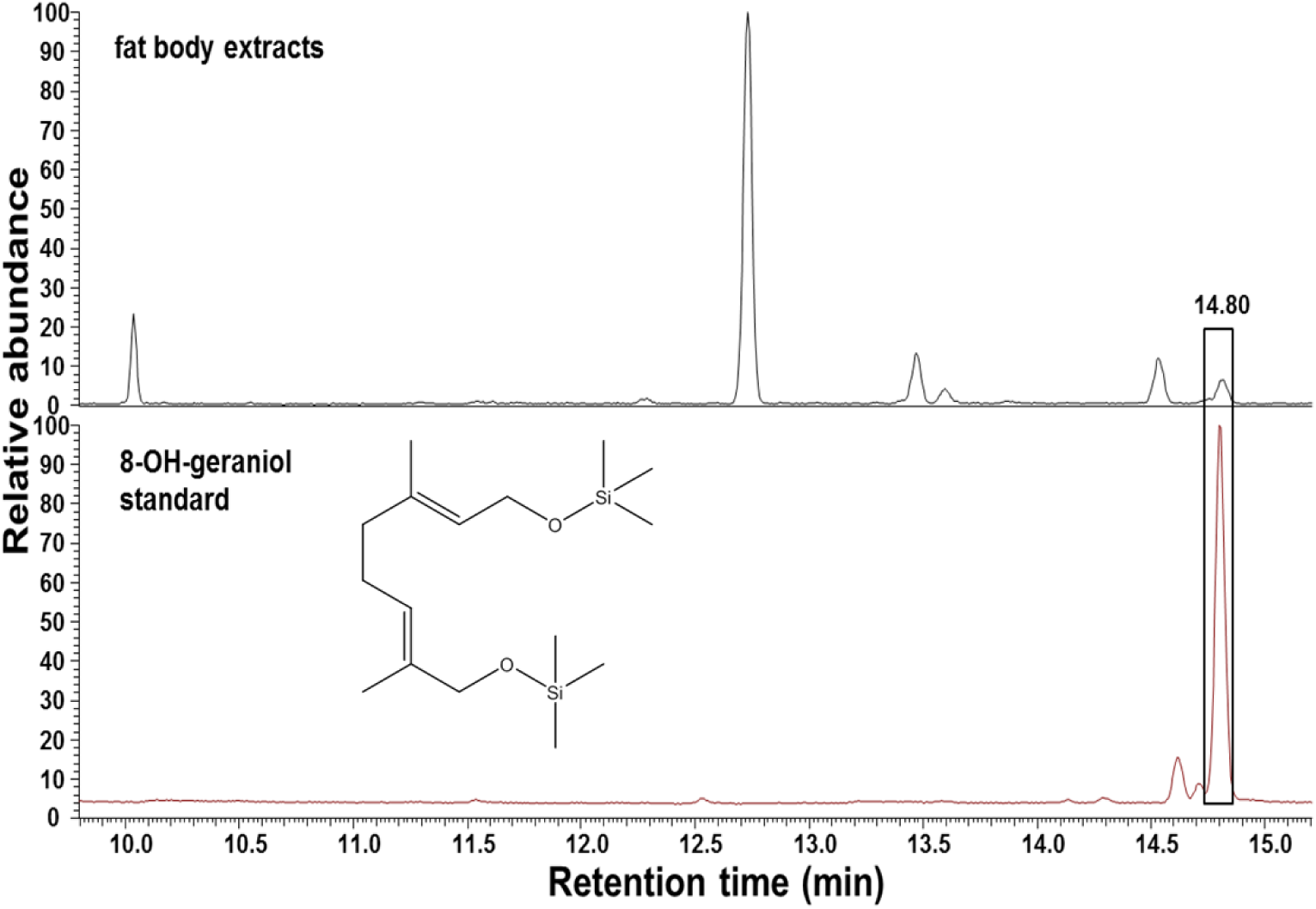
Identification of MSTFA silylated 8-OH-geraniol (RT: 14.80 min) in the fat body of *P. cochleariae* by GC-MS. 8-OH-geraniol in fat body extracts was assigned based on comparison of the retention time and mass spectra to the authentic standard.

### 3.2. Identification of CYP candidates from the *P. cochleariae* fat body

In order to identify the oxidase(s) involved in iridoid biosynthesis, approximately 193 transcripts (supplementary file 1) were annotated as putative cytochrome P450 in our in-house *P. cochleariae* transcriptomic reference library (Stock et al., 2013). However, based on amino acid identity to the homologous sequence of plant origin (Collu et al., 2001), none of them could be assigned as a dedicated geraniol 8-hydroxylase. To narrow down the number of potential P450s, the sequence information was combined with the proteomics analysis of the fat body membrane. A total of 73 cytochrome P450s were identified in the fat body, among them, 44 CYP6s, 12 CYP4s, 9 CYP9s, 5 CYP347s, and 1 each of CYP345, CYP 349 and CYP306 (Figure 3 and Table S1). Since the geraniol 8-hydroxylase was supposed to be a relatively abundant protein in the fat body, the matched peptide numbers of these candidates were also compared (Table S1). We selected 12 candidates that had more than 20 detected matched peptides for further analysis.

**Figure 3.**
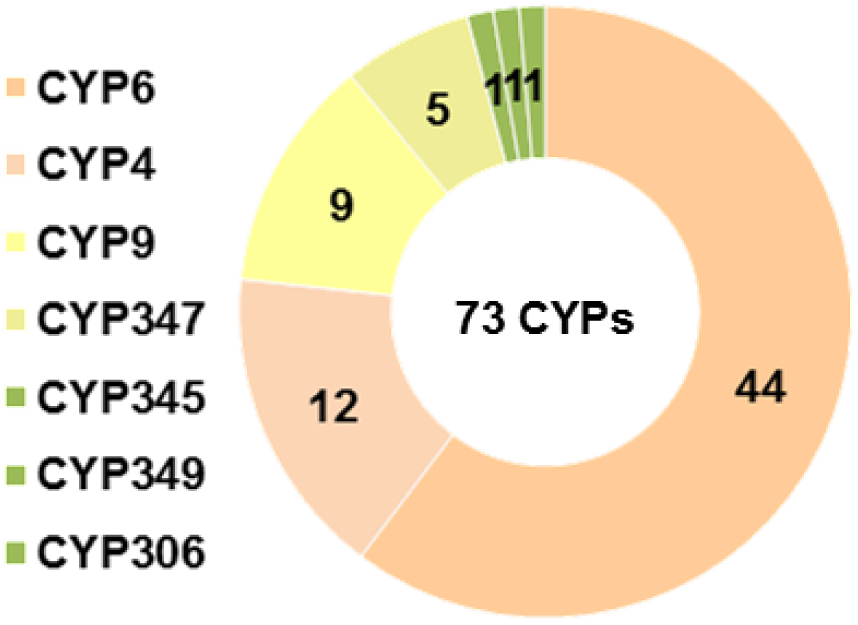
Cytochrome P450 numbers and families identified by membrane proteomics in the fat body. Different families were clustered by blasting against known CYPs in NCBI. The values represent the corresponding number of CYPs in each family.

The expression pattern of selected candidates was an additional criterion for identifying the prominent P450 members in *P. cochleariae* fat body. RT-qPCR was carried out to measure the relative abundance of the selected transcripts from different larval tissues, including the gut, Malpighian tubules, fat body and glands. Since *Pc* C7558, *Pc* C7614 and *Pc* C28218 differed from each other by only 1 or 2 amino acids (Fig. S3), therefore, *Pc C7558* was taken as a representative gene for RT-qPCR analysis. As shown in Figure 4 (Table S2), 9 genes were more abundantly expressed in the fat body than in the other tissues, whereas *Pc C16689* showed higher expression levels in the gut and Malpighian tubules than in the fat body. Thus, these 9 genes were considered as putative candidates for further RNAi analysis.

**Figure 4.**
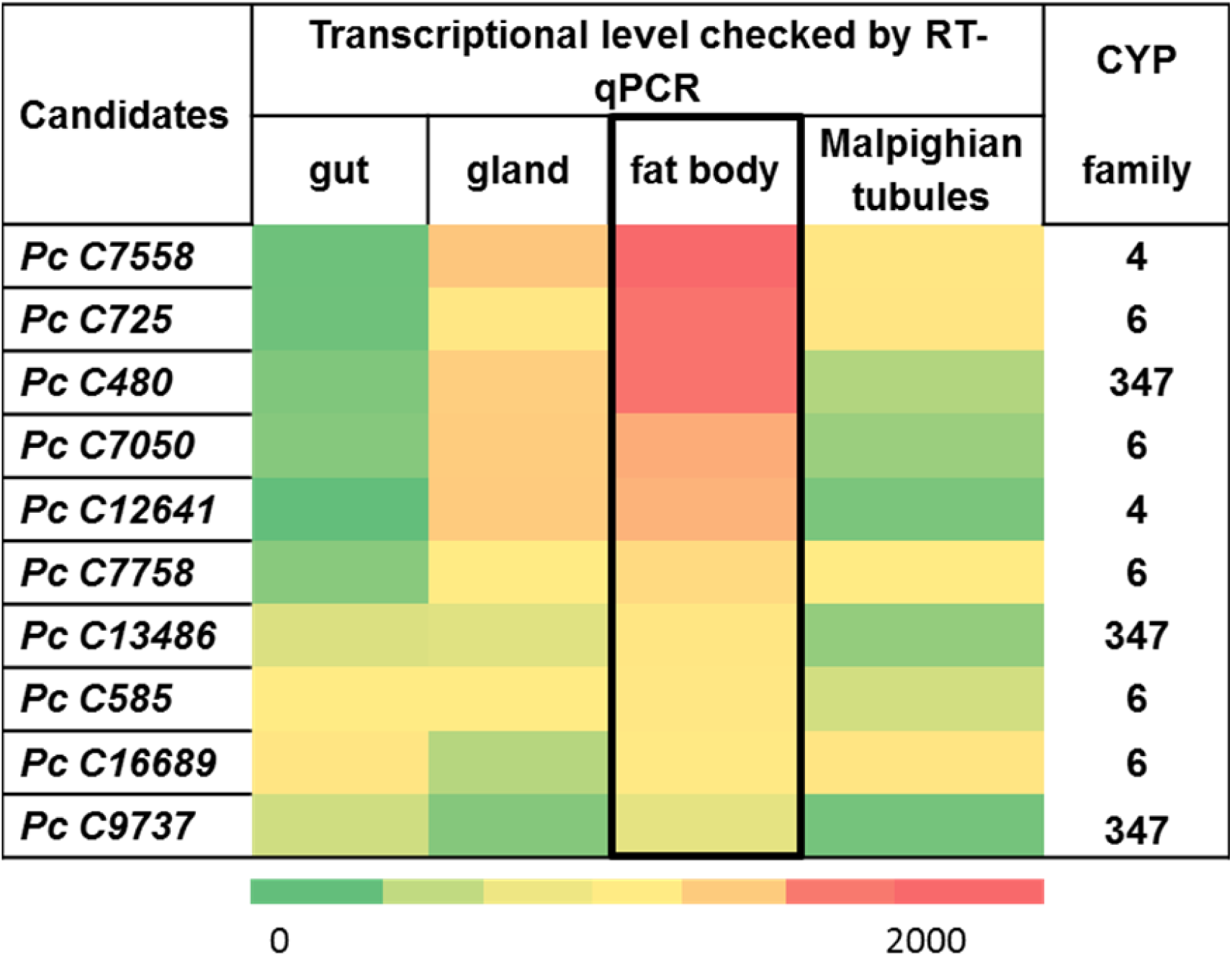
The tissue specific expression profile of the selected P450 candidates in *P. cochleariae* larvae. The heat map was generated based on the values measured by RT-qPCR. The data shown are mean + SE (n = 3). The color gradients from green to red represent the values ranging from 0 to 2000. For the detailed values, see Table S3. Pc *eIF4a* and Pc *EF1a* were used for normalizing the transcript abundance.

### 3.3. RNAi-based identification of P450(s) involved in iridoid biosynthesis

To pinpoint the right candidate for the iridoid biosynthesis, the dsRNA of the 9 genes was synthesized. We ranked and tested the individual candidates according to its transcripts abundance in the fat body, from most to least. Second instar *P. cochleariae* larvae were used for injection, and larvae treated with dsRNA targeting *eGFP* were viewed as a control group. In order to check whether the induced RNAi effect would influence iridoid biosynthesis, the amount of the downstream product 8-OH-Ger-Glc present in larval hemolymph was examined via HPLC-MS. Among the tested candidates, only when knocking down *Pc C7758* did we detect a significant decrease of 8-OH-Ger-Glc compared to the level in the *eGFP* control group (Fig. S4). We then further verified the observed effect caused by the suppression of *Pc C7758* by carrying out time course sampling. As indicated in Figure 5a, in the ds *eGFP* injected group, the relative 8-OH-Ger-Glc amount increased gradually from 2 to 7 days after injection and then remained at a relatively stable level. In the ds *C7758* injected group, in contrast, we observed a significant decrease on the fifth day after injection; and afterwards the compound remained at a basal low level until the larvae reached the pre-pupal stage.

**Figure 5.**
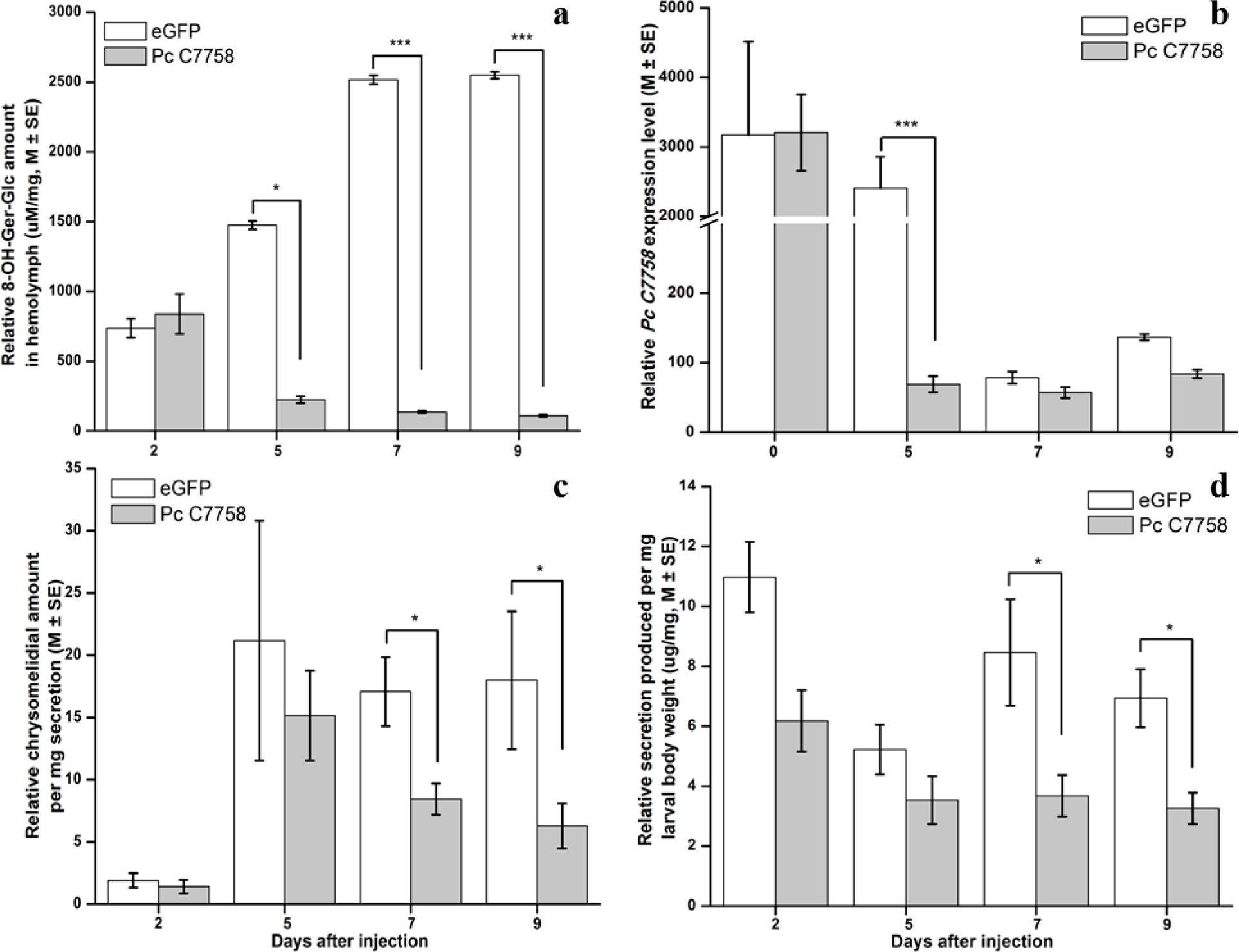
Timeline of different RNAi effects. a) Relative 8-hydroxygeraniol glucoside (8-OH-Ger-Glc) amount in hemolymph after RNAi (n≥9); b) Transcriptional level of *Pc C7758* (n = 3, each time point contains 3 biological replicates); c) Relative chrysomelidial amount in the glandular reservoir (n≥5); d) Relative glandular secretion produced by larvae in Pc C7758 knockdown and eGFP groups (n≥5, mean ± SE).

The observed decrease of 8-OH-Ger-Glc due to the down-regulation of *Pc C7758* was correlated with transcription analysis and larval physiological development. As shown in Figure 5b, the relative transcription level of *Pc C7758* in both groups showed identical and high expression upon injection. However, on the fifth day after injection, the transcriptional level in *Pc C7758* knockdown group dropped to 2.8% of that in the eGFP control group, indicating that the RNAi effect had been triggered and the gene had been successfully silenced. The observed low transcription level of *Pc* C7758 in the eGFP control from the seventh day on is likely due to the physiological down-regulation, as the larvae have reached the pre-pupal state, which is closely related to the non-chrysomelidial-containing pupal stage (Pasteels et al., 1988). Moreover, knocking down the expression of the *Pc* C7758 gene did not affect larval physiological development, as indicated by the similar body weight observed in the two experiment groups (Fig. S5).

In addition, we examined the knock down effect of *Pc C7758* on the larvae’s ability to produce defensive secretion and chrysomelidial (Fig. 5c-d). As shown in Figure 5c, the chrysomelidial concentration in the glandular secretion increased along with the body weight gain until it rose to the stable level that characterized earlier larval stages. Although, in the *Pc* C7758 knock down group, the chrysomelidial concentration reached peak level on the fifth day after injection, it decreased steadily and significantly afterwards due to the lack of *Pc C7758* transcripts. Figure 5d indicated that the larvae tended to maintain a relatively stable level of glandular secretion despite the continual body weight gain during the experimental period. However, the relative level in the *Pc* C7758 knock down group was always slightly less, and the difference became significant seven days after injection. Based on our results, we concluded that *P. cochleariae* recruit the cytochrome P450 *Pc* C7758 for iridoid biosynthesis. According to the P450 nomenclature committee, this sequence is referred to as CYP6BH5.

### 3.4. *In vitro* evidence of geraniol hydroxylase activity of CYP6BH5 and its substrate selectivity

Direct evidence of CYP6BH5’s function as a geraniol hydroxylase was obtained *in vitro* by the functional co-expression of CYP6BH5 and CPR derived from *C. populi*. A pilot experiment with the microsomal fraction from HEK 293 cells expressing CYP6BH5 and CPR showed the detectable transformation of geraniol to 8-OH-geraniol (Fig. S6). Due to the low yield of the recombinant proteins in HEK 293 cells, we produced CYP6BH5 and CPR in Sf9 insect cells via transient baculovirus-mediated expression (Fig. S7). Enzyme assays with recombinant CYP6BH5/CPR microsomes derived from Sf9 cells also revealed the conversion of geraniol into 8-OH-geraniol in the presence of NADPH. There was no detectable turnover of the substrate in either negative-control reaction (without NADPH) or empty control reaction (microsome containing only CPR), see Figure 6A (mass spectra, see Fig. S1). Because members from the CYP76 family (such as CYP76B6) were recently reported as representing versatile monoterpenol oxidases (Boachon et al., 2015; Hofer et al., 2014), a set of monoterpenols that are structurally related to geraniol, namely nerol, linalool and citronellol, were additionally tested. No conversion was detected when linalool was used as a substrate. However, the hydroxylation of nerol and citronellol by CYP6BH5 was observed. NMR data verified that nerol was converted to 8-hydroxynerol (Fig. 6B, mass spectra and NMR data, see Fig. S8 and S10a). Similarly, citronellol was metabolized to 8-hydroxycitronellol (Fig. 6C, mass spectra and NMR data, see Fig. S9 and S10b). Sung et al. (2011) showed CYP76B6 from *C. roseus* also plays a role in phenylpropanoid biosynthesis, catalyzing 3-hydroxylation of naringenin to produce eriodictyol. Thus, we also test CYP6BH5’s activity on naringenin. However, no reaction product was detected compared to the negative or empty control.

**Figure 6.**
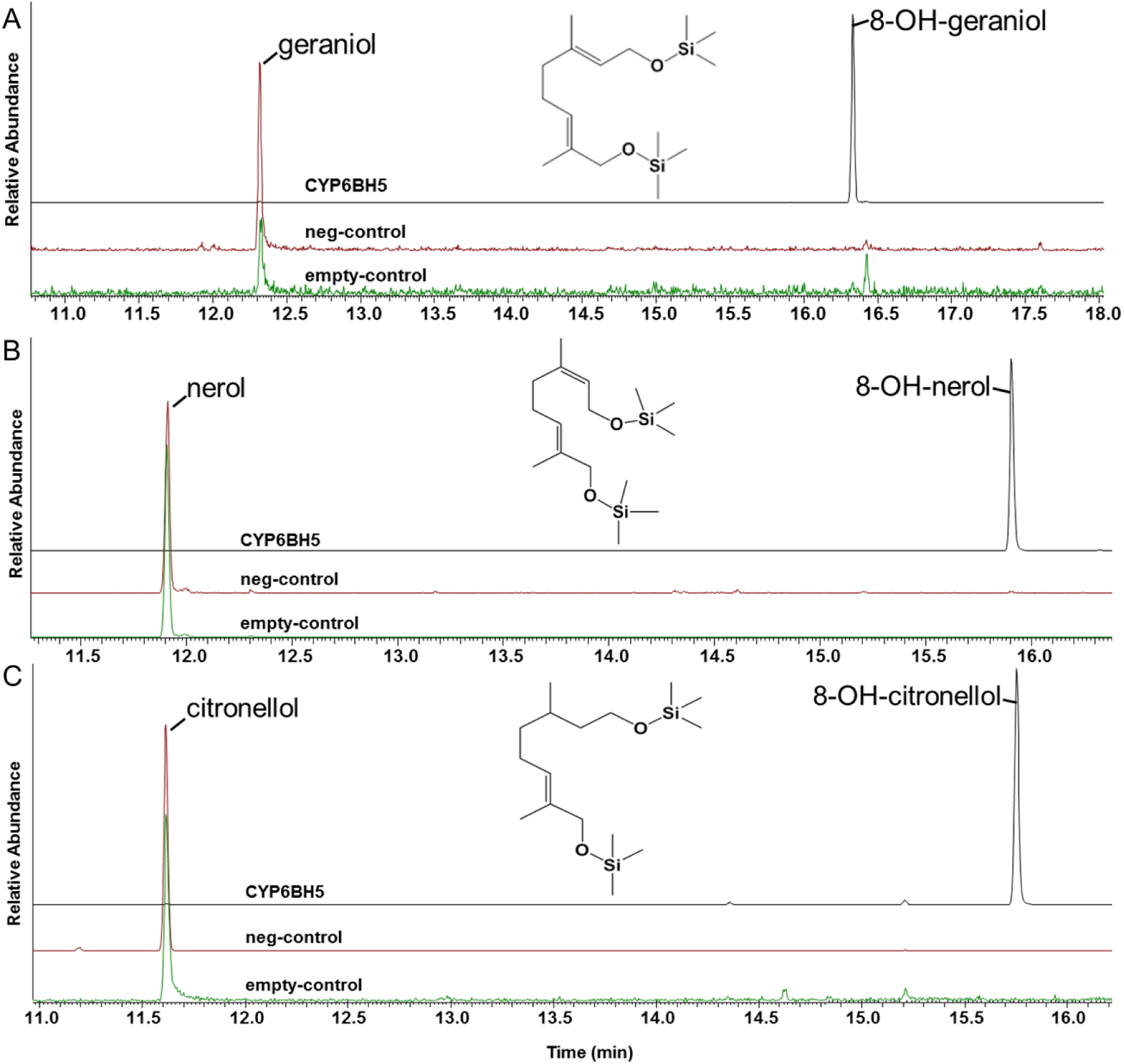
GC chromatograms of the reaction products show the conversion of monoterpenols by the CYP6BH5 enzymes expressed by Sf9 insect cells. Microsomal membranes from Sf9 insect cells co-express CYP6BH5 and CPR or of CPR only (empty-control) were incubated with 200 µM of substrate for 15 min in the presence of NADPH. No NADPH was added to the negative control (neg-control). Identified compounds were then assigned based on their retention times and MS data. Mass spectra and NMR data of the products and references are available in supplementary Figures S1 and S6 to S10.

### 3.5 Homology modeling of CYP6BH5 with the substrate geraniol

To reveal the underlying mechanism of CYP6BH5’s substrate preference for geraniol (1-ol) over linalool (3-ol), the docking of geraniol to a homology-based 3D-structure of CYP6BH5 was conducted. As shown in Figure 7, geraniol was positioned close to the hydroperoxyl heme of the enzyme’s active site. The docking pose is stabilized by the formation of hydrogen bonds between the hydroxyl group of geraniol and the side chain of Arg205, a carboxyl group of the heme prosthetic group and with Thr369. The hydrophobic side chain of Val365 also helps to stabilize the correct position by hydrophobic interactions with the prenyl moiety of geraniol. The transoid methyl group of geraniol has a distance of 3.2 Å, compared to 3.4 Å of the cisoid methyl group, to the distal oxygen atom sitting in the middle of the heme center; this explained why CYP6BH5 preferentially leads to the oxidation of the *trans* methyl group of the substrate. Since the substrate geraniol was positioned by a hydrogen bond between its 1-OH group and Arg205, it is reasonable to assume that the embedding of citronellol and nerol into the active center of CYP6BH5 is controlled by the same forces leading to ω-hydroxylation at their *trans*-methyl groups. The 3-OH group of linalool led to a wrong positioning and, hence, this compound was not oxidized.

**Figure 7.**
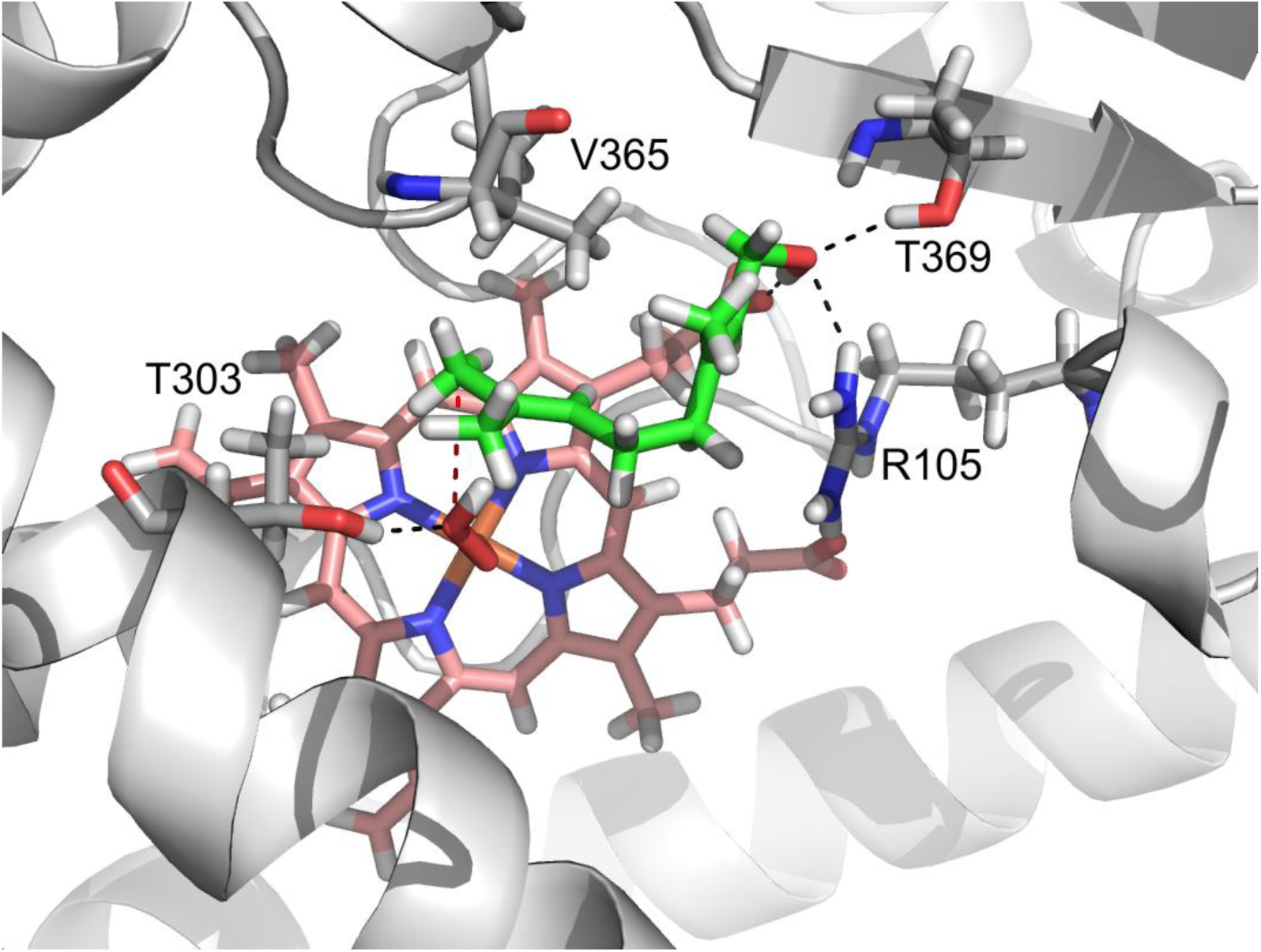
Docking arrangement of geraniol in the active site of CYP6BH5. The hydroxyl group of geraniol is recognized by hydrogen bonds with T369 and Arg105 as well as with the carboxyl moiety of heme. Furthermore, the side chain of Val365 supports the correct positioning by hydrophobic interactions with the prenyl moiety. The transoid methyl group has a distance of 3.2 Å and the cisoid methyl group of 3.4 Å to the distal oxygen atom sitting in the middle of the heme center. Therefore, the enzyme will preferentially lead to oxidation of the *trans* methyl group. Oxygen is red, carbon is green, nitrogen is blue, hydrogen is white and iron is brown.

## 4. Discussion

The cytochrome P450s (P450s) comprise a superfamily of heme-containing monooxygenases that are known to catalyze a broad range of endogenous substrates as well as xenobiotics in all living organisms (Scott, 1999). The abundance of P450s in insects varies greatly from species to species, although P450s in insects have not undergone the degree of gene duplication seen in higher plants (Schuler, 2011). In this study, we identified 73 P450s in the fat body of *P. cochleariae* larvae by a combination of proteomic and transcriptomic analyse. Based on *in vivo* and *in vitro* assays, our data convincingly proved that the *P. cochleariae* CYP6BH5 is an essential enzyme for the production of the defensive monoterpenoid chrysomelidial, by converting geraniol to 8-OH-geraniol.

Among the transcripts of the 73 annotated *P. cochleariae* fat body P450s, more than 60% transcripts (44) were allocated to the CYP6 family by comparing these with the known insect P450s in NCBI. In general, members from the CYP6 family are well known for mediating the ecological adaptation of herbivorous insects to their host plants (Schuler, 2011). In addition, they have also been reported to help the insects develop resistance by detoxifying plant secondary metabolites or pesticides (Scott, 1999). The best-characterized xenobiotic metabolizing CYP6Bs are ascribed to members isolated from Lepidoptera species (Li et al., 2001, 2004; Mao et al., 2007; Wen et al., 2006). For example, in the corn earworm *Helicoverpa zea*, a generalist feeding on hundreds of host plant species, multiple CYP6B subfamily members co-exist (Li et al., 2002b). Among them, CYP6B8 has been reported to metabolize a broad range of plant allelochemicals including flavone, xanthotoxin, chlorogenic acid, indole-3-carbinol, quercetin, and rutin (Li et al., 2004). Additionally, the same enzyme can also metabolize three different insecticides such as cypermethrin, diazinon and aldrin (Hung et al., 1996; Li et al., 2004, 2002a; Rupasinghe et al., 2007; Schuler, 2011). In *P. cochleariae*, 31 out of the 44 annotated putative fat body CYP6 transcripts were clustered into the CYP6B subfamily. The occurrence of abundant and diverse CYP6B transcripts in the *P. cochleariae* fat body implicates their potential roles in xenobiotic metabolism.

Out of all the CYP6B transcripts identified in this study, only CYP6BH5 could be correlated with iridoid biosynthesis in *P. cochleariae*. Beyond geraniol, *in vitro* enzyme assays showed CYP6BH5 metabolizes other monoterpene alcohols, such as nerol and citronellol, but not linalool. Homology modeling of CYP6BH5 with geraniol showed the –OH group of the substrate played an important role in coordinating the substrate with the enzyme’s catalytic cavity. Compared to geraniol, linalool differs from its isomer in terms of its –OH position, with –OH located at C3 for linalool, at C1 for geraniol. This difference prevents the prenyl moiety of linalool from becoming close enough to the heme center of CYP6BH5; therefore, linalool could not be hydroxylated. However, Boachon et al. (2015) showed *P. cochleariae* could feed on linalool-releasing Arabidopsis as well as the cabbage leaves treated with pure linalool. Hence, it is postulated that *P. cochleariae* harbors a presumable P450 that differs from the geraniol 8-hydroxylase CYP6BH5 for linalool metabolism.

In *P. cochleariae*, geraniol is catalyzed to 8-OH-geraniol by CYP6BH5 for iridoid biosynthesis. In plants, geraniol is known as a constituent component of essential oils in more than 160 species (especially the *Cymbopogon* genus) and exhibits insecticidal and repellent properties in response to insects such as mosquitoes, cockroaches ticks, etcetera (Burdock, 2010; Chen and Viljoen, 2010). Therefore, it is hypothesized that other insects may also have enzymes to detoxify geraniol. How common is it for members from the CYP6B subfamily in other insects to be recruited to metabolize geraniol and even monoterpene alcohols metabolism? To answer this question, more enzymatic information is needed. Until now, CYP6BH5 is the first identified insect P450 that metabolizes geraniol and its structurally related monoterpene alcohols, nerol and citronellol. In plants, members from the CYP76C subfamily and some other CYP76s, CYP76B6 for example, are also closely associated with monoterpenol oxidation (Collu et al., 2001; Hofer et al., 2013). However, compared to these plant-derived P450s, CYP6BH5 showed a narrower substrate spectrum, as indicated by the absence of detectable activity on linalool and the phenylpropanoid pathway intermediate naringenin. This feature reveals that *P. cochleariae* may recruit CYP6BH5 specifically for geraniol hydroxylation in the iridoid biosynthesis pathway. In spite of that, CYP6BH5, like members from CYP76s, is a substrate-promiscuous enzyme with regard to its activity on monoterpenols (Hofer et al., 2013, 2014). The observed substrate spectrum similarity among CYPs that are not closely related is likely to result from convergent evolution.

Apart from *P. cochleariae*, the Madagascar periwinkle (*C. roseus*) can also use geraniol as precursor for iridoid biosynthesis (Miettinen et al., 2014). In this plant, the iridoid (nepetalactol) serves as a precursor for secologanin, which can be metabolized further into pharmacologically important monoterpenoid indole alkaloids, such as vinblastine and vincristine (Alagna et al., 2016; Caputi et al., 2018; Kellner et al., 2015; Miettinen et al., 2014). In both organisms, the biosynthetic processes from geraniol to iridoid share similar sequential steps, although not identical enzymes. Though both organisms recruit a member from the cytochrome P450 superfamily to generate 8-OH-geraniol, the geraniol 8-hydroxylase CYP6BH5 from *P. cochleariae* shares only 20% amino acid identity to CYP76B6 in *C. roseus*. Afterwards, *P. cochleariae* uses a 8-hydroxygeraniol oxidase (*Pc* 8HGO) from the GMC oxidoreductases superfamily to produce 8-oxogeranial, whereas *C. roseus* takes either the CYP76B6 or a NAD+-dependent 8-hydroxygeraniol oxidoreductase belonging to the medium-chain reductase superfamily to catalyze this reaction (Hofer et al., 2013; Miettinen et al., 2014; Rahfeld et al., 2014). For the ring closure step, a member of the Rossmann-fold NAD(P)+-binding protein superfamily, namely, the iridoid synthase was used to cyclize the 8-oxogeranial in *C. roseus* (Geu-Flores et al., 2012). In *P. cochleariae*, although knocking down a member of the juvenile hormone-binding protein superfamily leads to the accumulation of the precursor 8-oxogeranial in glandular secretion, the heterologously expressed protein showed no activity on 8-oxogeranial (Bodemann et al., 2012).

The elucidation of the involved enzymes in the iridoid biosynthesis pathway in both *P. cochleariae* and *C. roseus* lays the foundation for producing these pharmacologically valuable monoterpenoid indole alkaloids on an industrial level. Recent, attempts to reconstruct the biosynthetic pathways for nepetalactol, vindoline and strictosidine from the geranyl diphosphate in yeast *Saccharomyces cerevisiae* have made impressive progress (Billingsley et al., 2017; Brown et al., 2015; Campbell et al., 2016; Qu et al., 2015). Despite the effort to use genetic engineering to optimize the performance of the platform strains, the ω-hydroxylation of geraniol at C-8 remains a bottleneck (Brown et al., 2015). The newly identified CYP6BH5 from *P. cochleariae*, therefore, may serve as an alternative biocatalyst for overcoming this obstacle, further boosting the yields of monoterpenoid indole alkaloids.

## Acknowledgments

We sincerely thank Dr. Maritta Kunert for her support in chemical analysis, Dr. Ding Wang and Lydia Schmidt for their help with bioinformatics, Prof. Stefan H. Heinemann for his help with HEK cells, and Dr. Nicole Joußen for her constructive suggestions about P450 enzymes. The authors would also like to express their gratitude to Sarah Baur for her help in constructing the plasmids, to David Nelson (University of Tennessee) for help with P450 nomenclature and to Emily Wheeler for her critical reading of the manuscript. This work was financially supported by the Max Planck Society and the China Scholarship Council [grant number 201406300098].

## Reference

Alagna, F., Geu-Flores, F., Kries, H., Panara, F., Baldoni, L., O’Connor, S.E., Osbourn, A., 2016. Identification and characterization of the iridoid synthase involved in oleuropein biosynthesis in olive (Olea europaea) fruits. J Biol Chem 291, 5542–5554. https://doi.org/10.1074/jbc.M115.701276

Berman, H.M., Westbrook, J., Feng, Z., Gilliland, G., Bhat, T.N., Weissig, H., Shindyalov, I.N., Bourne, P.E., 2000. The protein Ddata bank. Nucleic Acids Res 28, 235–242. http://www.rcsb.org/pdb/

Billingsley, J.M., DeNicola, A.B., Barber, J.S., Tang, M.-C., Horecka, J., Chu, A., Garg, N.K., Tang, Y., 2017. Engineering the biocatalytic selectivity of iridoid production in Saccharomyces cerevisiae. Metab. Eng. 44, 117–125. https://doi.org/10.1016/j.ymben.2017.09.006

Boachon, B., Junker, R.R., Miesch, L., Bassard, J.-E., Höfer, R., Caillieaudeaux, R., Seidel, D.E., Lesot, A., Heinrich, C., Ginglinger, J.-F., Allouche, L., Vincent, B., Wahyuni, D.S.C., Paetz, C., Beran, F., Miesch, M., Schneider, B., Leiss, K., Werck-Reichhart, D., 2015. CYP76C1 (Cytochrome P450)-Mediated linalool metabolism and the formation of volatile and soluble linalool oxides in Arabidopsis flowers: A strategy for defense against floral antagonists. Plant Cell 27, 2972. https://doi.org/10.1105/tpc.15.00399%0A

Bodemann, R.R., Rahfeld, P., Stock, M., Kunert, M., Wielsch, N., Groth, M., Frick, S., Boland, W., Burse, A., 2012. Precise RNAi-mediated silencing of metabolically active proteins in the defence secretions of juvenile leaf beetles. Proc. R. Soc. B-Biological Sci. 279, 4126–4134. https://doi.org/10.1098/rspb.2012.1342

Boros, C.A., Stermitz, F.R., 1991. Iridoids-An updated review.2. J. Nat. Prod. 54, 1173–1246. https://doi.org/10.1021/np50077a001

Brown, S., Clastre, M., Courdavault, V., O’Connor, S.E., 2015. De novo production of the plant-derived alkaloid strictosidine in yeast. Proc Natl Acad Sci U S A 112, 3205–3210. https://doi.org/10.1073/pnas.1423555112

Burdock, G., 2010. Fenaroli’s Handbook of Flavor Ingredients., 6th Edition. ed, Boca Raton: CRC Press. https://doi.org/10.1201/9781439847503

Burse, A., Frick, S., Discher, S., Tolzin-Banasch, K., Kirsch, R., Strauss, A., Kunert, M., Boland, W., 2009. Always being well prepared for defense: The production of deterrents by juvenile Chrysomelina beetles (Chrysomelidae). Phytochemistry 70, 1899–1909. https://doi.org/10.1016/j.phytochem.2009.08.002

Burse, A., Schmidt, A., Frick, S., Kuhn, J., Gershenzon, J., Boland, W., 2007. Iridoid biosynthesis in Chrysomelina larvae: Fat body produces early terpenoid precursors. Insect Biochem. Mol. Biol. 37, 255–265. https://doi.org/10.1016/j.ibmb.2006.11.011

Bustin, S.A., Benes, V., Garson, J.A., Hellemans, J., Huggett, J., Kubista, M., Mueller, R., Nolan, T., Pfaffl, M.W., Shipley, G.L., Vandesompele, J., Wittwer, C.T., 2009. The MIQE guidelines: minimum information for publication of quantitative real-time PCR experiments. Clin Chem 55, 611–622. https://doi.org/10.1373/clinchem.2008.112797

Campbell, A., Bauchart, P., Gold, N.D., Zhu, Y., De Luca, V., Martin, V.J., 2016. Engineering of a nepetalactol-producing platform strain of Saccharomyces cerevisiae for the production of plant seco-iridoids. ACS Synth Biol 5, 405–414. https://doi.org/10.1021/acssynbio.5b00289

Caputi, L., Franke, J., Farrow, S.C., Chung, K., Payne, R.M.E., Nguyen, T.-D., Dang, T.-T.T., Soares Teto Carqueijeiro, I., Koudounas, K., Dugé de Bernonville, T., Ameyaw, B., Jones, D.M., Vieira, I.J.C., Courdavault, V., O’Connor, S.E., 2018. Missing enzymes in the biosynthesis of the anticancer drug vinblastine in Madagascar periwinkle. Science. 360(6394), 1235–1239. https://doi.org/10.1126/science.aat4100

Cavill, G.W.K., Robertson, P.L., Brophy, J.J., Duke, R.K., McDonald, J., Plant, W.D., 1984. Chemical ecology of the meat ant, Iridomyrmex purpureus sens. strict. Insect Biochem. 14, 505–513. https://doi.org/10.1016/0020-1790(84)90004-0

Chen, W., Viljoen, A.M., 2010. Geraniol - A review of a commercially important fragrance material. South African J. Bot. 76, 643–651. https://doi.org/10.1016/j.sajb.2010.05.008

Collu, G., Unver, N., Peltenburg-Looman, A.M.G., van der Heijden, R., Verpoorte, R., Memelink, J., 2001. Geraniol 10-hydroxylase, a cytochrome P450 enzyme involved in terpenoid indole alkaloid biosynthesis. Febs Lett. 508, 215–220. https://doi.org/10.1016/s0014-5793(01)03045-9

Frick, S., Nagel, R., Schmidt, A., Bodemann, R.R., Rahfeld, P., Pauls, G., Brandt, W., Gershenzon, J., Boland, W., Burse, and A., 2013. Metal ions control product specificity of isoprenyl diphosphate synthases in the insect terpenoid pathway. Proc Natl Acad Sci U S A 110(11):4194–4199. https://doi.org/10.1073/pnas.1221489110

Geu-Flores, F., Sherden, N.H., Courdavault, V., Burlat, V., Glenn, W.S., Wu, C., Nims, E., Cui, Y., O/’Connor, S.E., 2012. An alternative route to cyclic terpenes by reductive cyclization in iridoid biosynthesis. Nature 492, 138–142. https://doi.org/10.1038/nature11692

Ghisalberti, E.L., 1998. Biological and pharmacological activity of naturally occurring iridoids and secoiridoids. Phytomedicine 5, 147–163. https://doi.org/10.1016/S0944-7113(98)80012-3

Hofer, R., Boachon, B., Renault, H., Gavira, C., Miesch, L., Iglesias, J., Ginglinger, J.F., Allouche, L., Miesch, M., Grec, S., Larbat, R., Werck-Reichhart, D., 2014. Dual function of the cytochrome P450 CYP76 family from Arabidopsis thaliana in the metabolism of monoterpenols and phenylurea herbicides. Plant Physiol 166, 1149–1161. https://doi.org/10.1104/pp.114.244814

Hofer, R., Dong, L., Andre, F., Ginglinger, J.F., Lugan, R., Gavira, C., Grec, S., Lang, G., Memelink, J., Van der Krol, S., Bouwmeester, H., Werck-Reichhart, D., 2013. Geraniol hydroxylase and hydroxygeraniol oxidase activities of the CYP76 family of cytochrome P450 enzymes and potential for engineering the early steps of the (seco)iridoid pathway. Metab Eng 20, 221–232. https://doi.org/10.1016/j.ymben.2013.08.001

Hung, C.F., Holzmacher, R., Connolly, E., Berenbaum, M.R., Schuler, M.A., 1996. Conserved promoter elements in the CYP6B gene family suggest common ancestry for cytochrome P450 monooxygenases mediating furanocoumarin detoxification. Proc. Natl. Acad. Sci. U. S. A. 93, 12200–12205. https://doi.org/10.1073/pnas.93.22.12200

Joussen, N., Agnolet, S., Lorenz, S., Schone, S.E., Ellinger, R., Schneider, B., Heckel, D.G., 2012. Resistance of Australian Helicoverpa armigera to fenvalerate is due to the chimeric P450 enzyme CYP337B3. Proc Natl Acad Sci U S A 109, 15206–15211. https://doi.org/10.1073/pnas.1202047109

Kellner, F., Geu-Flores, F., Sherden, N.H., Brown, S., Foureau, E., Courdavault, V., O’Connor, S.E., 2015. Discovery of a P450-catalyzed step in vindoline biosynthesis: a link between the aspidosperma and eburnamine alkaloids. Chem. Commun. 51, 7626–7628. https://doi.org/10.1039/C5CC01309G

Krieger, E., Joo, K., Lee, J., Lee, J., Raman, S., Thompson, J., Tyka, M., Baker, D., Karplus, K., 2009. Improving physical realism, stereochemistry, and side-chain accuracy in homology modeling: Four approaches that performed well in CASP8. Proteins 77 Suppl 9, 114–122. https://doi.org/10.1002/prot.22570

Krieger, E., Vriend, G., 2014. YASARA View - molecular graphics for all devices - from smartphones to workstations. Bioinformatics 30, 2981–2982. https://doi.org/10.1093/bioinformatics/btu426

Krithika, R., Srivastava, P.L., Rani, B., Kolet, S.P., Chopade, M., Soniya, M., Thulasiram, H. V, 2015. Characterization of 10-Hydroxygeraniol dehydrogenase from Catharanthus roseus reveals cascaded enzymatic activity in iridoid biosynthesis. Sci. Rep. 5, 6. https://doi.org/10.1038/srep08258

Kunert, M., Rahfeld, P., Shaker, K.H., Schneider, B., David, A., Dettner, K., Pasteels, J.M., Boland, W., 2013. Beetles do it differently: Two stereodivergent cyclisation modes in iridoid-producing leaf-beetle larvae. Chembiochem 14, 353–360. https://doi.org/10.1002/cbic.201200689

Kunert, M., Soe, A., Bartram, S., Discher, S., Tolzin-Banasch, K., Nie, L., David, A., Pasteels, J., Boland, W., 2008. De novo biosynthesis versus sequestration: A network of transport systems supports in iridoid producing leaf beetle larvae both modes of defense. Insect Biochem. Mol. Biol. 38, 895–904. https://doi.org/10.1016/j.ibmb.2008.06.005

Larsen, B., Fuller, V.L., Pollier, J., Van Moerkercke, A., Schweizer, F., Payne, R., Colinas, M., O’Connor, S.E., Goossens, A., Halkier, B.A., 2017. Identification of iridoid glucoside transporters in Catharanthus roseus. Plant Cell Physiol 58, 1507–1518. https://doi.org/10.1093/pcp/pcx097

Laskowski, R.A., MacArthur, M.W., Moss, D.S., Thornton, J.M., 1993. ProcHECK: a program to check the stereochemical quality of protein structures. J. Appl. Crystallogr. 26, 283–291. https://doi.org/doi:10.1107/S0021889892009944

Laurent, P., Braekman, J.-C., Daloze, D., 2005. Insect Chemical Defense, in: Schulz, S. (Ed.), The Chemistry of Pheromones and Other Semiochemicals II. Springer Berlin Heidelberg, Berlin, Heidelberg, pp. 167–229. https://doi.org/10.1007/b98317

Li, W., Berenbaum, M.R., Schuler, M.A., 2001. Molecular analysis of multiple CYP6B genes from polyphagous Papilio species. Insect Biochem Mol Biol 31, 999–1011. https://doi.org/10.1016/S0965-1748(01)00048-0

Li, X., Baudry, J., Berenbaum, M.R., Schuler, M.A., 2004. Structural and functional divergence of insect CYP6B proteins: From specialist to generalist cytochrome P450. Proc Natl Acad Sci U S A 101, 2939–2944. https://doi.org/10.1073/pnas.0308691101

Li, X., Berenbaum, M.R., Schuler, M.A., 2002a. Cytochrome P450 and actin genes expressed in Helicoverpa zea and Helicoverpa armigera: paralogy/orthology identification, gene conversion and evolution. Insect Biochem Mol Biol 32, 311–320. https://doi.org/10.1016/S0965-1748(01)00092-3

Li, X., Schuler, M.A., Berenbaum, M.R., 2002b. Jasmonate and salicylate induce expression of herbivore cytochrome P450 genes. Nature 419, 712–715. https://doi.org/10.1038/nature01003

Mao, W., Schuler, M.A., Berenbaum, M.R., 2007. Cytochrome P450s in Papilio multicaudatus and the transition from oligophagy to polyphagy in the Papilionidae. Insect Mol Biol 16, 481–490. https://doi.org/10.1111/j.1365-2583.2007.00741.x

Miettinen, K., Dong, L., Navrot, N., Schneider, T., Burlat, V., Pollier, J., Woittiez, L., van der Krol, S., Lugan, R., Ilc, T., Verpoorte, R., Oksman-Caldentey, K.-M., Martinoia, E., Bouwmeester, H., Goossens, A., Memelink, J., Werck-Reichhart, D., 2014. The seco-iridoid pathway from Catharanthus roseus. Nat Commun 5. https://doi.org/10.1038/ncomms4606

Oldham, N.J., Veith, M., Boland, W., Dettner, K., 1996. Iridoid monoterpene biosynthesis in insects: Evidence for a de novo pathway occurring in the defensive glands of Phaedon armoraciae (Chrysomelidae) leaf beetle larvae. Naturwissenschaften 83, 470–473. https://doi.org/10.1007/BF01144016

Pasteels, J., Braekman, J.-C., Daloze, D., n.d. Chemical defense in the Chrysomelidae, in: Jolivet, P., Petitpierre, E., Hsiao, T.H. (Eds.), Biology of Chrysomelidae, Series Entomologica. Springer Netherlands, pp. 233–252. https://doi.org/10.1007/978-94-009-3105-3_14

Qu, Y., Easson, M.L.A.E., Froese, J., Simionescu, R., Hudlicky, T., De Luca, V., 2015. Completion of the seven-step pathway from tabersonine to the anticancer drug precursor vindoline and its assembly in yeast. Proc. Natl. Acad. Sci. 112, 6224–6229. https://doi.org/10.1073/pnas.1501821112

Rahfeld, P., Haeger, W., Kirsch, R., Pauls, G., Becker, T., Schulze, E., Wielsch, N., Wang, D., Groth, M., Brandt, W., Boland, W., Burse, A., 2015. Glandular beta-glucosidases in juvenile Chrysomelina leaf beetles support the evolution of a host-plant-dependent chemical defense. Insect Biochem. Mol. Biol. 58, 28–38. https://doi.org/10.1016/j.ibmb.2015.01.003

Rahfeld, P., Kirsch, R., Kugel, S., Wielsch, N., Stock, M., Groth, M., Boland, W., Burse, A., 2014. Independently recruited oxidases from the glucose-methanol-choline oxidoreductase family enabled chemical defences in leaf beetle larvae (subtribe Chrysomelina) to evolve. Proc. R. Soc. B-Biological Sci. 281. https://doi.org/10.1098/rspb.2014.0842

Rosa, T., Monica, R.L., Federica, M., Giancarlo, A.S., Francesco, M., 2008. Biological and Pharmacological activities of iridoids: Recent developments. Mini-Reviews Med. Chem. 8, 399–420. http://dx.doi.org/10.2174/138955708783955926

Rupasinghe, S.G., Wen, Z., Chiu, T.L., Schuler, M.A., 2007. Helicoverpa zea CYP6B8 and CYP321A1: different molecular solutions to the problem of metabolizing plant toxins and insecticides. Protein Eng Des Sel 20, 615–624. https://doi.org/10.1093/protein/gzm063

Schuler, M.A., 2011. P450s in plant-insect interactions. Biochim. Biophys. Acta-Proteins Proteomics 1814, 36–45. https://doi.org/10.1016/j.bbapap.2010.09.012

Scott, J.G., 1999. Cytochromes P450 and insecticide resistance. Insect Biochem. Mol. Biol. 29, 757–777. https://doi.org/10.1016/s0965-1748(99)00038-7

Sippl, M.J., 1993. Recognition of errors in three-dimensional structures of proteins. Proteins 17, 355–362. https://doi.org/10.1002/prot.340170404

Sippl, M.J., 1990. Calculation of conformational ensembles from potentials of mean force. An approach to the knowledge-based prediction of local structures in globular proteins. J Mol Biol 213, 859–883. https://doi.org/10.1016/S0022-2836(05)80269-4

Smith, R.M., Brophy, J.J., Cavill, G.W.K., Davies, N.W., 1979. Iridodials and nepetalactone in the defensive secretion of the coconut stick insects,Graeffea crouani. J. Chem. Ecol. 5, 727–735. https://doi.org/10.1007/BF00986557

Snyder, J.H., Qi, X., 2013. Biosynthesis: Metal matters. Nat Chem Biol 9, 295–296. https://doi.org/10.1038/nchembio.1232

Stock, M., Gretscher, R.R., Groth, M., Eiserloh, S., Boland, W., Burse, A., 2013. Putative sugar transporters of the mustard leaf beetle Phaedon cochleariae: their phylogeny and role for nutrient supply in larval defensive glands. PLoS One 8, e84461. https://doi.org/10.1371/journal.pone.0084461

Strauss, A.S., Peters, S., Boland, W., Burse, A., 2013. ABC transporter functions as a pacemaker for sequestration of plant glucosides in leaf beetles. Elife 2, e01096. http://dx.doi.org/10.7554/eLife.01096.001

Sung, P.H., Huang, F.C., Do, Y.Y., Huang, P.L., 2011. Functional expression of geraniol 10-hydroxylase reveals its dual function in the biosynthesis of terpenoid and phenylpropanoid. J. Agric. Food Chem. 59, 4637–4643. https://doi.org/10.1021/jf200259n

Veith, M., Lorenz, M., Boland, W., Simon, H., Dettner, K., 1994. Biosynthesis of iridoid monoterpenes in insects: Defensive secretions from larvae of leaf beetles (coleoptera: chrysomelidae). Tetrahedron 50, 6859–6874. http://dx.doi.org/10.1016/S0040-4020(01)81338-7

Wen, Z., Rupasinghe, S., Niu, G., Berenbaum, M.R., Schuler, M.A., 2006. CYP6B1 and CYP6B3 of the black swallowtail (Papilio polyxenes): adaptive evolution through subfunctionalization. Mol Biol Evol 23, 2434–2443. https://doi.org/10.1093/molbev/msl118

Yamane, H., Konno, K., Sabelis, M., Takabayashi, J., Sassa, T., Oikawa, H., 2010. Chemical defence and toxins of plants, in: Comprehensive natural products II. Elsevier, Oxford, pp. 339–385. https://doi.org/10.1016/B978-008045382-8.00099-X

