## Supplementary Figures for "A cytochrome P450 from juvenile mustard leaf beetles hydroxylates geraniol, a key step in iridoid biosynthesis"

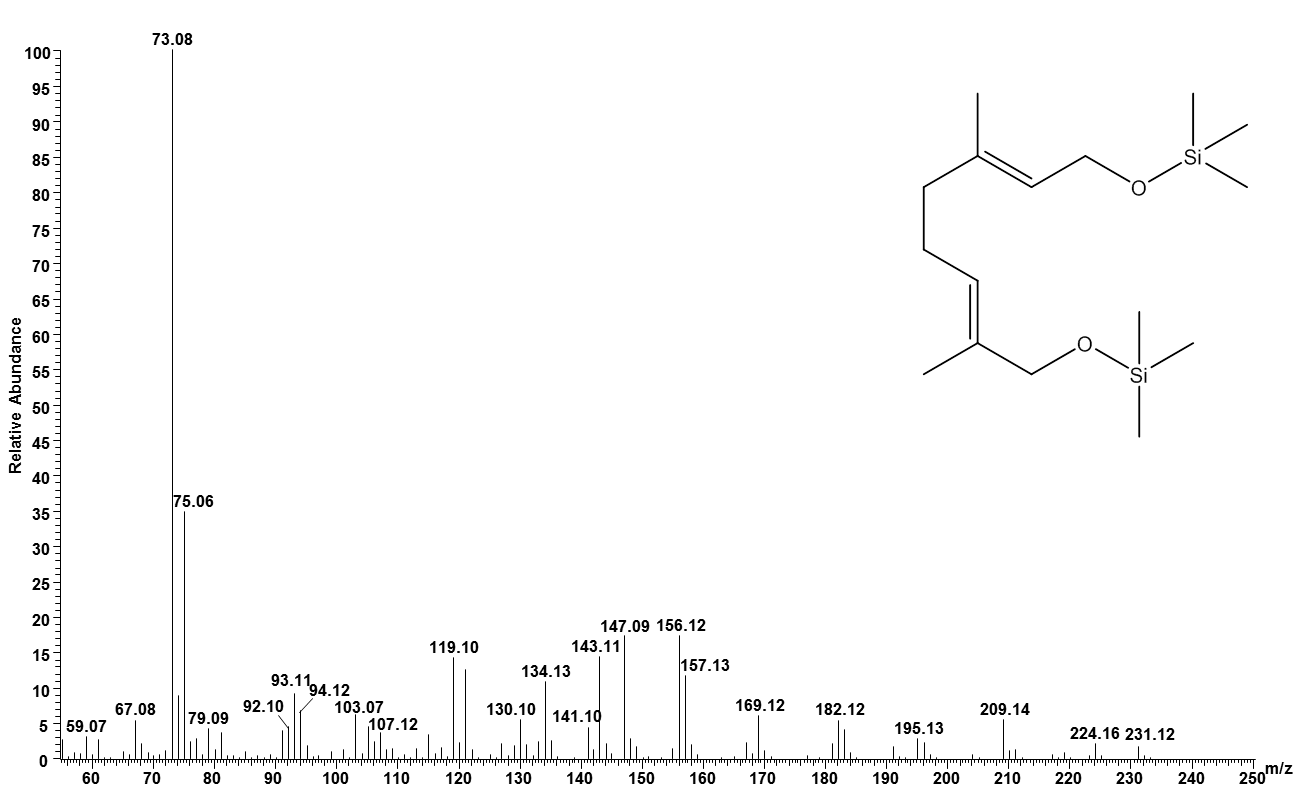

Fig S1 EI-MS of MSTFA silylated 8-OH-geraniol.

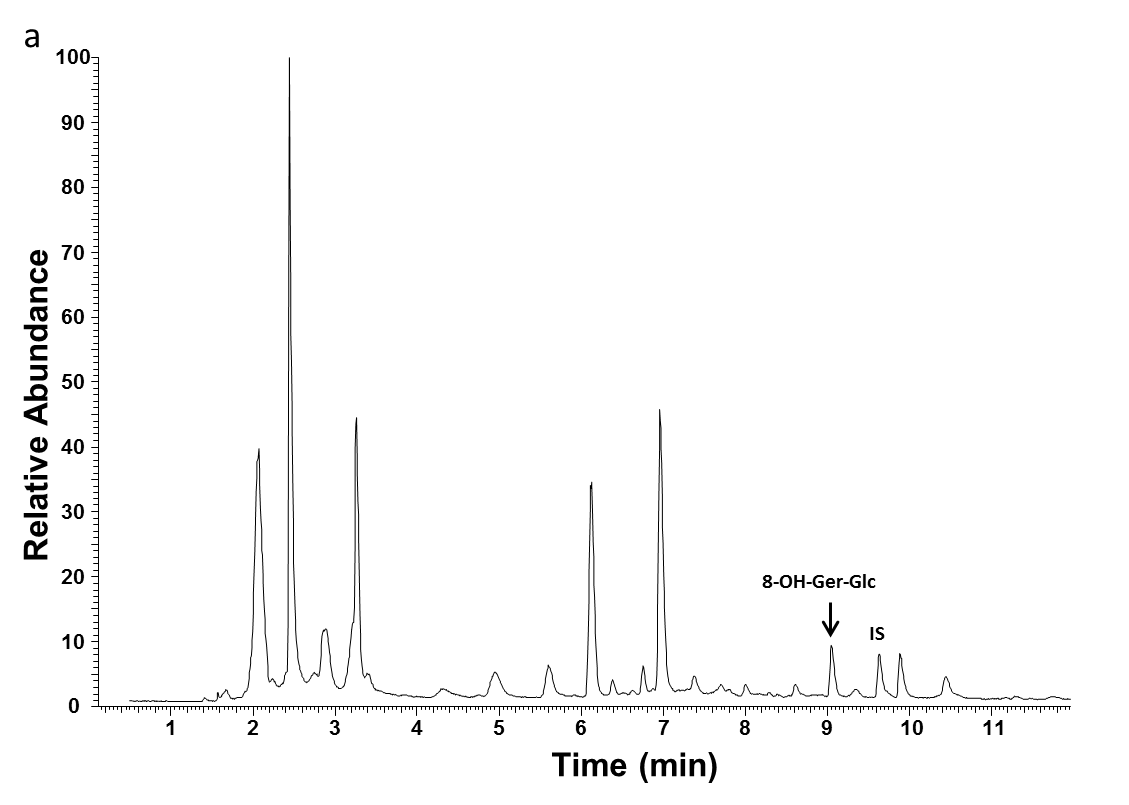

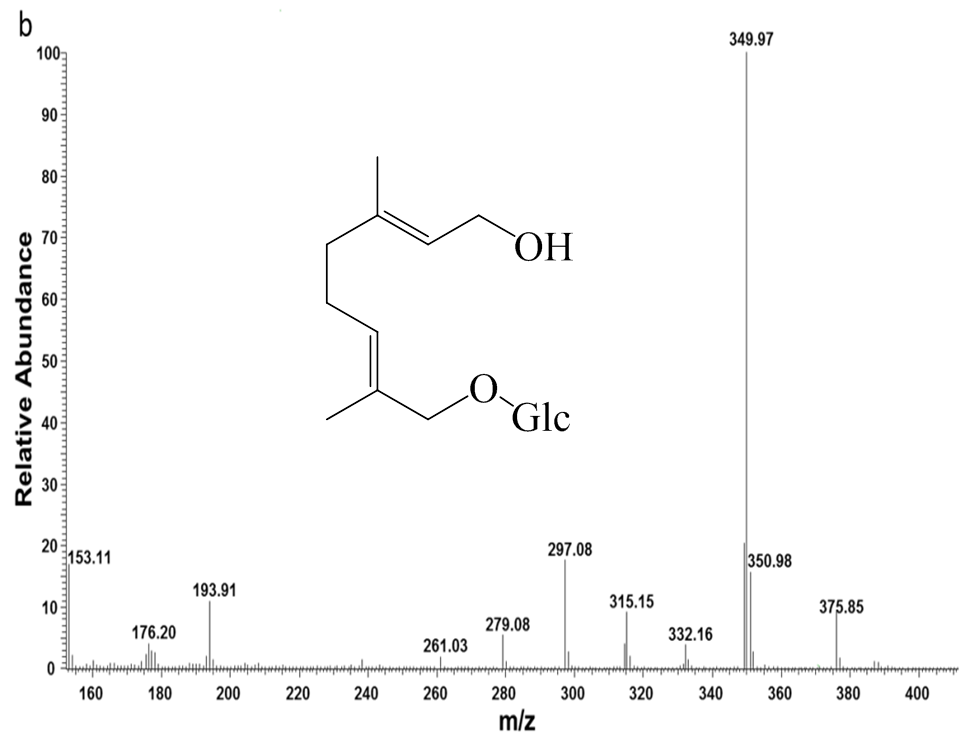

**1**

**2**

**3**

**4**

**5**

**6**

**7**

**8**

**9**

**10**

**11**

**Time (min)**

**0**

**10**

**20**

**30**

**40**

**50**

**60**

**70**

**80**

**90**

**100**

**Relative Abundance**

**IS**

**8-OH-Ger-Glc**

Fig S2 Identification of 8-hydroxygeraniol-8-O-*β*-D-glucoside (RT: 9.04 min) in the fat body of *P. cochleariae* by HPLC-APCI-MS. The thioglucoside of 8-hydroxygeraniol served as internal standard (IS, 9.62 min). a. Liquid chromatograph of *P. cochleariae* fat body extracts; b. mass spectra of 8-hydroxygeraniol-8-O-*β*-D-glucoside.

```
PcC7558     1 MSTTTASPDLAAPSALLSASSVFYFLLIPAAILWYAYWKISRSHMLELANKIPGPPGLPI
PcC28218    1 MSTTTASPDLAAPSALLSASSVFYFLLIPAAILWYAYWKISRSHMLELANKIPGPPGLPI
PcC7614     1 MSTTTASPDLAAPSALLSASSVFYFLLIPAAILWYAYWKISRSHMLELANKIPGPPGLPI


PcC7558    61 LGNALQLLGSSPQIFKRVYELSYDYGNTVKMWAGPKLIIFLIDPSDVEIILSSHVHIDKA
PcC28218   61 LGNALQLLGSSPQIFKRVYELSYDYGNTVKMWAGPKLIIFLIDPSDVEIILSSHVHIDKA
PcC7614    61 LGNALQLLGSSPQIFKRVYELSYDYGNTVKMWAGPKLIIFLIDPSDVEIILSSHVHIDKA


PcC7558   121 SEYRFFKPWLGDGLLISTGQKWRAHRKLIAPTFHLNVLKSFIDLFNANSREVVQKLKKEV
PcC28218  121 SEYRFFKPWLGDGLLISTGQKWRAHRKLIAPTFHLNVLKSFIDLFNANSREVVQKLKKEV
PcC7614   121 SEYRFFKPWLGDGLLISTGQKWRAHRKLIAPTFHLNVLKSFIDLFNANSREVVQKLKKEV


PcC7558   181 GKEFDCHDYMSEATVEILLETAMGVSKKTQDQSGYDYAKAVMDMCDILHLRHTKIWLRPD
PcC28218  181 GKEFDCHDYMSEATVEILLETAMGVSKKTQDQSGYDYAKAVMDMCDILHLRHTKIWLRPD
PcC7614   181 GKEFDCHDYMSEATVEILLETAMGVSKKTQDQSGYDYAKAVMDMCDILHLRHTKIWLRPD


PcC7558   241 IIFNFTKYAKVQEGLINVIHSLTRKVIKRKRADFEKGIRGSTAEVPEEAKTNSVGSVASK
PcC28218  241 IIFNFTRYAKVQEGLINVIHSLTRKVIKRKRADFEKGIRGSTAEVPEEAKTNSVGSVASK
PcC7614   241 IIFNFTRYAKVQEGLINVIHSLTRKVIKRKRADFEKGIRGSTAEVPEEAKTNSVGSVASK


PcC7558   301 TVVEGLSYGQSVGLTDDLDVDDDIGEKKRMAFLDLMIEASQNGVVINDEEIKEQVDTIMF
PcC28218  301 TVVEGLSYGQSVGLTDDLDVDDDIGEKKRMAFLDLMIEASQNGVVINDEEIKEQVDTIMF
PcC7614   301 TVVEGLSYGQSVGLTDDLDVDDDIGEKKRMAFLDLMIEASQNGVVINDEEIKEQVDTIMF


PcC7558   361 EGHDTTAAGSSFFLSMMGIHQDIQDKVIQEIDEIFGDSDRPATFADTLEMKYLERCMMET
PcC28218  361 EGHDTTAAGSSFFLSMMGIHQDIQDKVIQEIDEIFGDSDRPATFADTLEMKYLERCMMET
PcC7614   361 EGHDTTAAGSSFFLSMMGIHQDIQDKVIQEIDEIFGDSDRPATFADTLEMKYLERCMMET


PcC7558   421 LRMYPPVPIIARQLRQDVKLVSGDYTLPAGATIVIGTFKIHRLPDIYPNPDKFDPDNFLP
PcC28218  421 LRMYPPVPIIARQLRQDVKLVSGDYTLPAGATIVIGTFKIHRLPDIYPNPDKFDPDNFLP
PcC7614   421 FRMYPPVPIIARQLRQDVKLVSGDYTLPAGATIVIGTFKIHRLPDIYPNPDKFDPDNFLP


PcC7558   481 ERTANRHYYSFIPFSAGPRSCVGRKYAMLKLKILLSTILRNYRVRSDIQEKDFQLQADII
PcC28218  481 ERTANRHYYSFIPFSAGPRSCVGRKYAMLKLKILLSTILRNYRVRSDIQEKDFQLQADII
PcC7614   481 ERTANRHYYSFIPFSAGPRSCVGRKYAMLKLKILLSTILRNYRVRSDIQEKDFQLQADII


PcC7558   541 LKRAEGFKIKLEPRKRLAAAM
PcC28218  541 LKRAEGFKIKLEPRKRLAAAM
PcC7614   541 LKRAEGFKIKLEPRKRLAAAM
```

Fig S3 Amino acid sequence alignment of PcC7558, PcC28218 and PcC7614 (the different amino acids were marked in red).

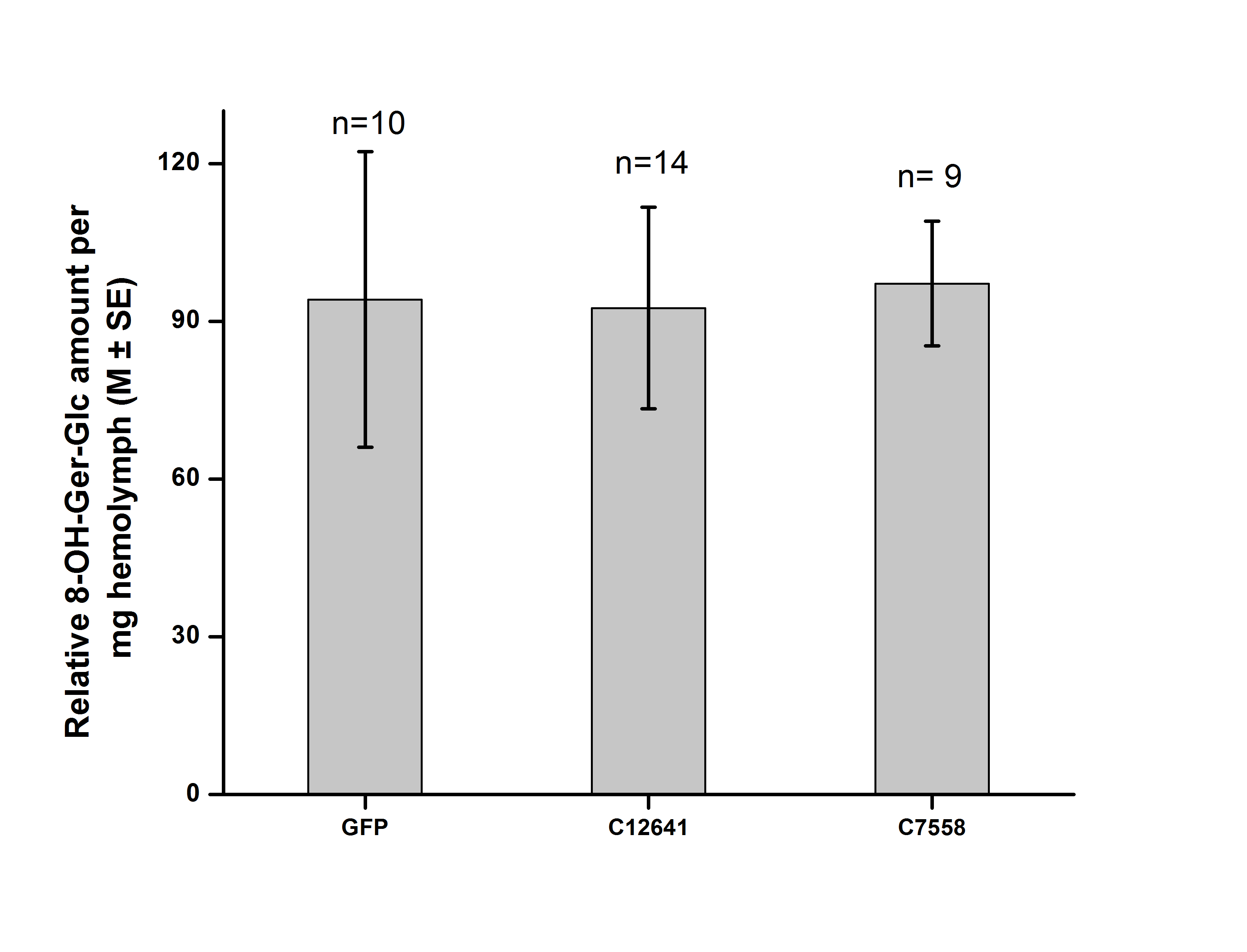

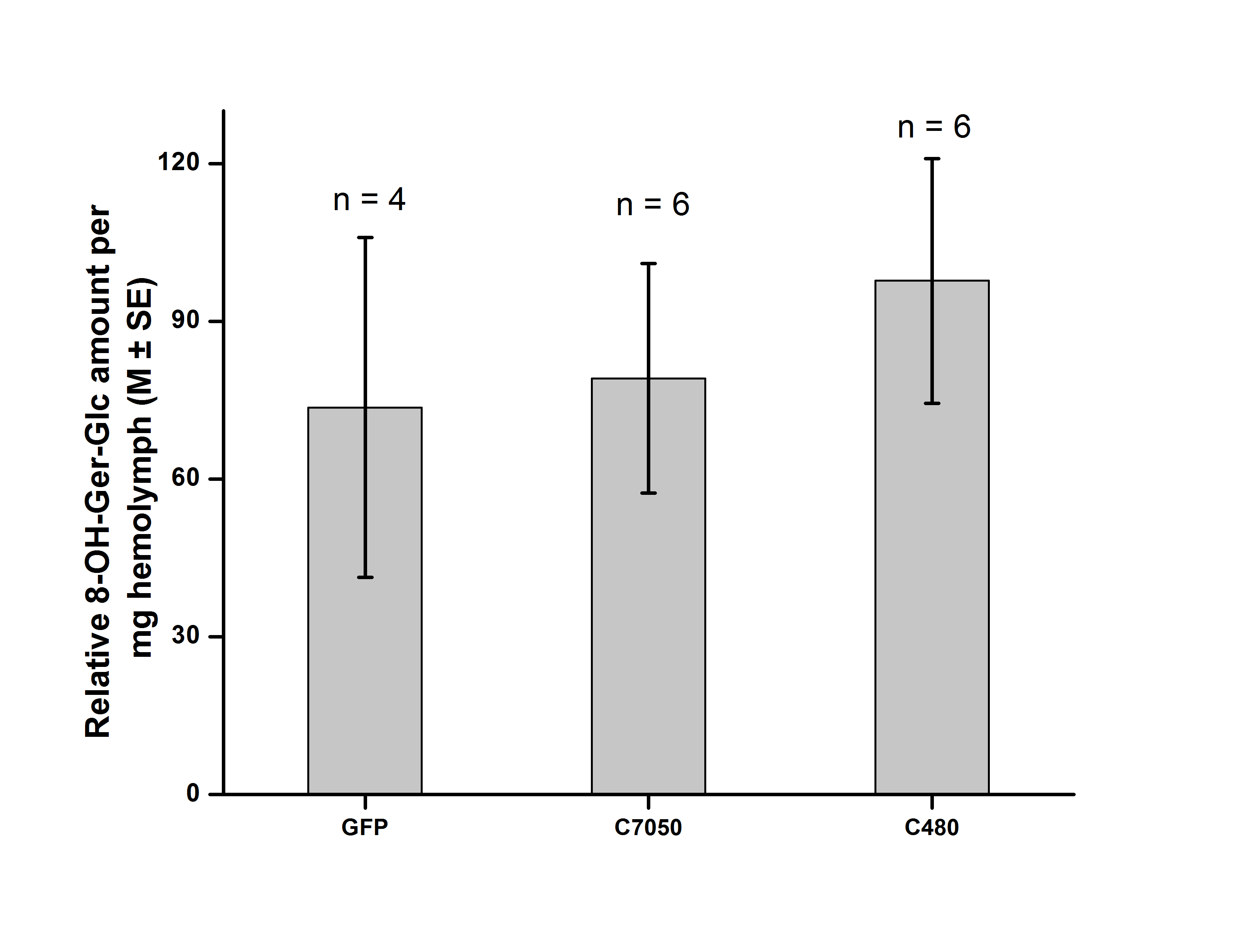

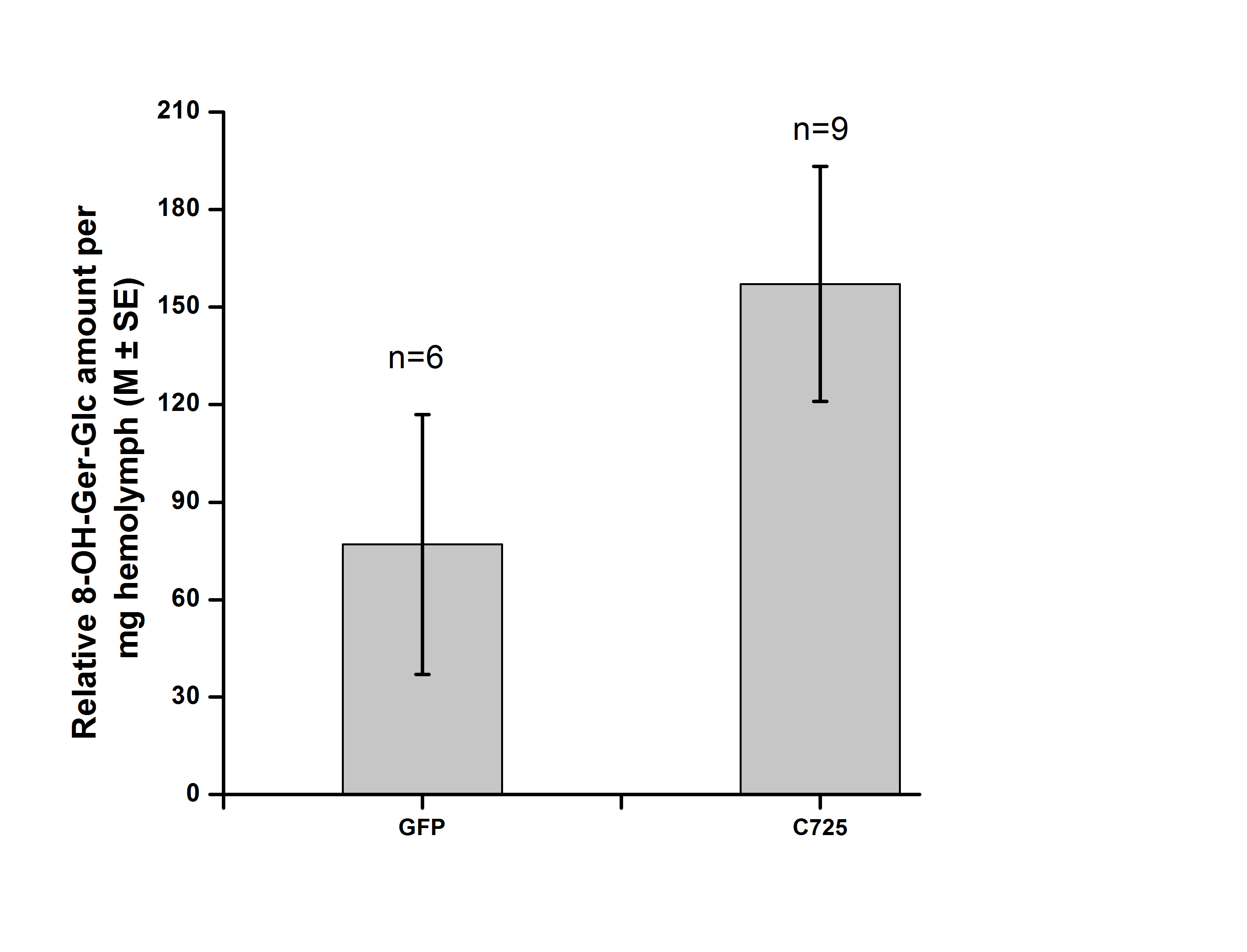

Fig S4 Relative 8-hydroxygeraniol glucosides amount in hemolymph on 7th day after dsRNA injection (M±SE)

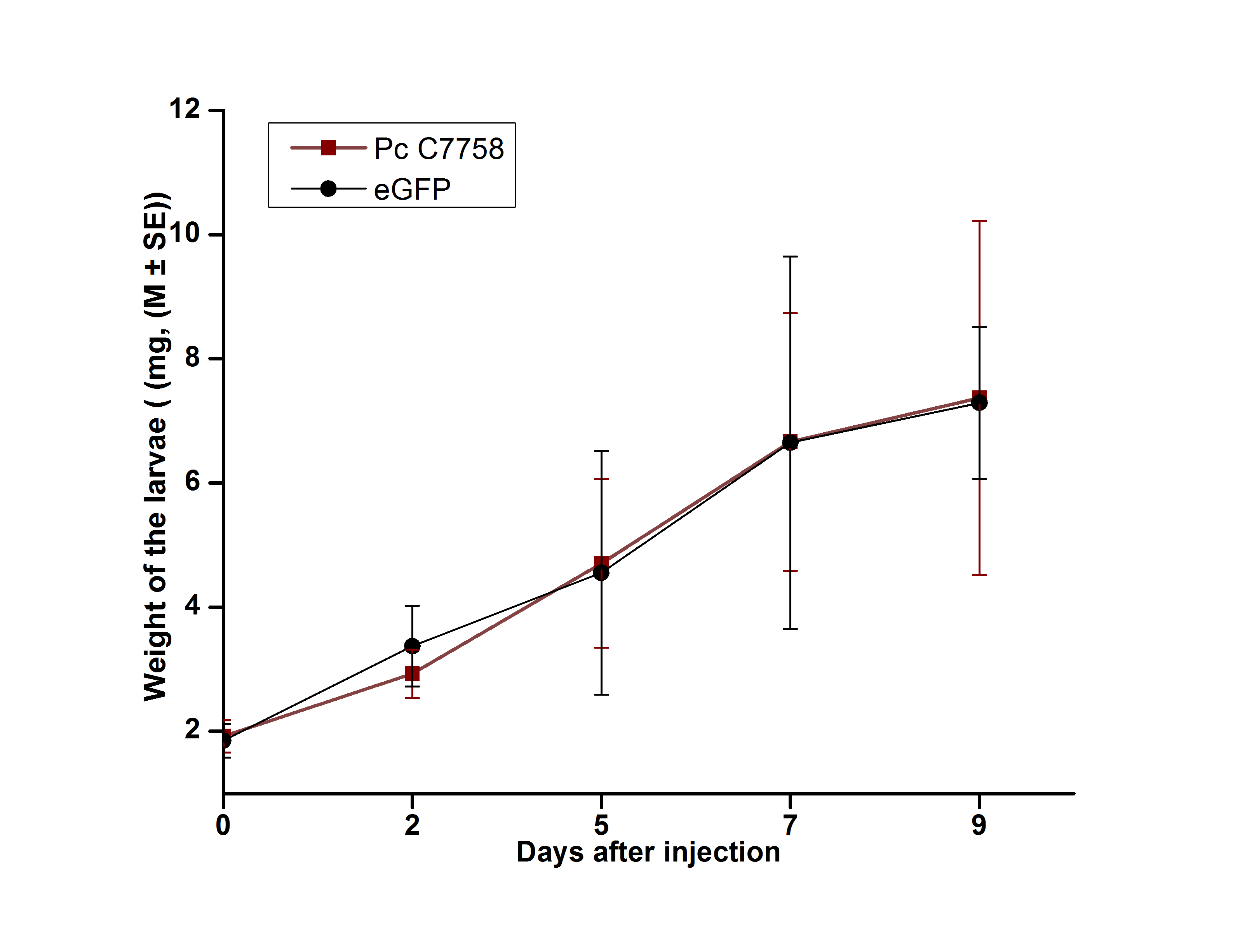

Fig S5 *P. cochleriae* larval fitness (body mass) was not influenced by knockdown of *Pc* *C7758* (n ≥ 9).

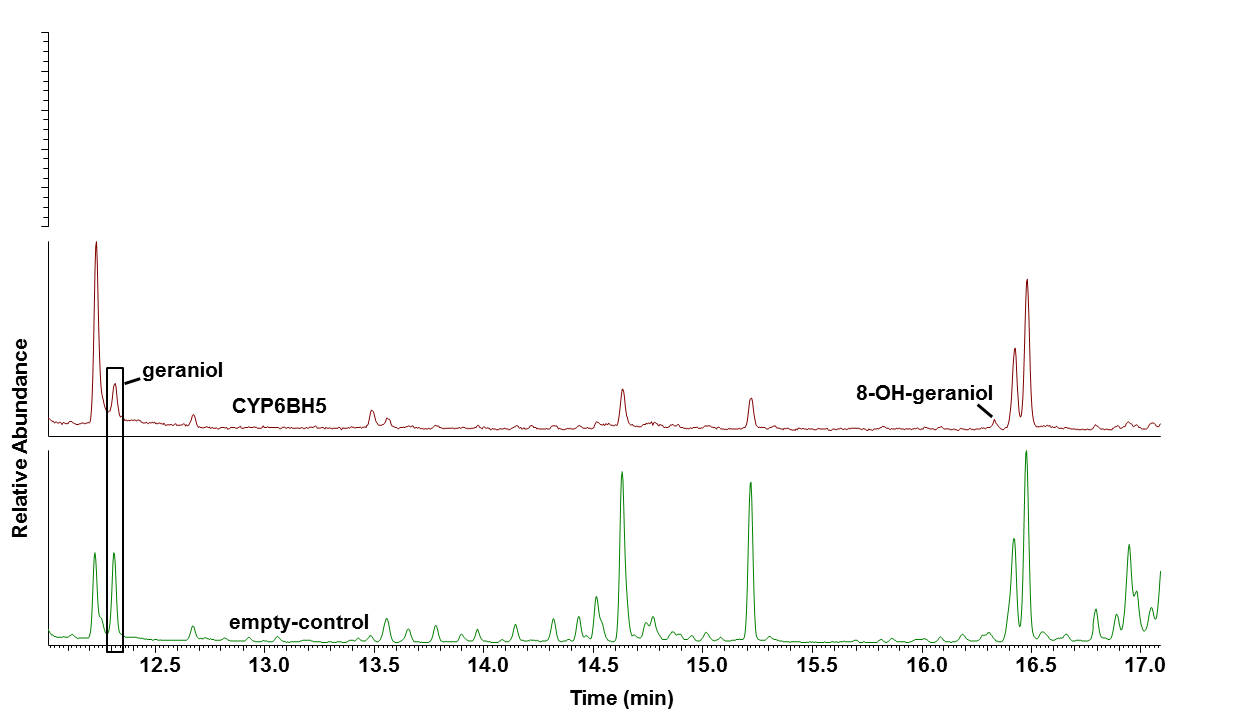

Fig S6 GC chromatograms of the reaction product results from the conversion of geraniol by the HEK293-expressed CYP6BH5 enzymes. Microsomal membranes from HEK293 cells co-transfected with vectors containing CYP6BH5 and CPR or of CPR only (empty-control) were incubated with 200 mM of substrate for 1 hour in the presence of NADPH.

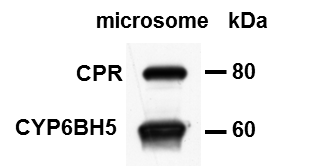

Fig S7 Western blot detection using Anti-V5-HRP antibody. Microsome was prepared from Sf9 cells that were co-transfected with recombinant baculoviral vectors expressing CYP6BH5 and CPR that fused to V5 epitope and His-tag. The observed molecular weights agree with predictions from their sequences.

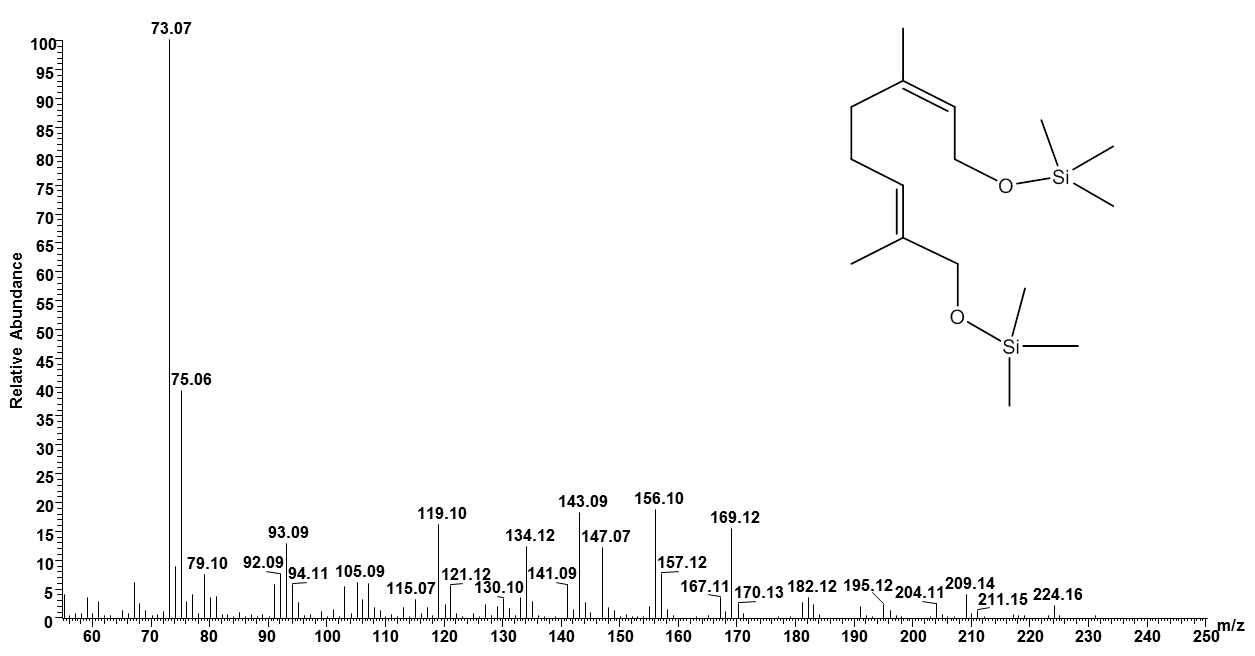

Fig S8: EI-MS of MSTFA silylated 8-hydroxynerol. EI mass spectrum corresponding to the peak of 8-hydroxynerol formed by CYP6BH5 in Figure 6B. According to NMR data acquired from the non-silylated compound, the peak is identical with 8-hydroxynerol.

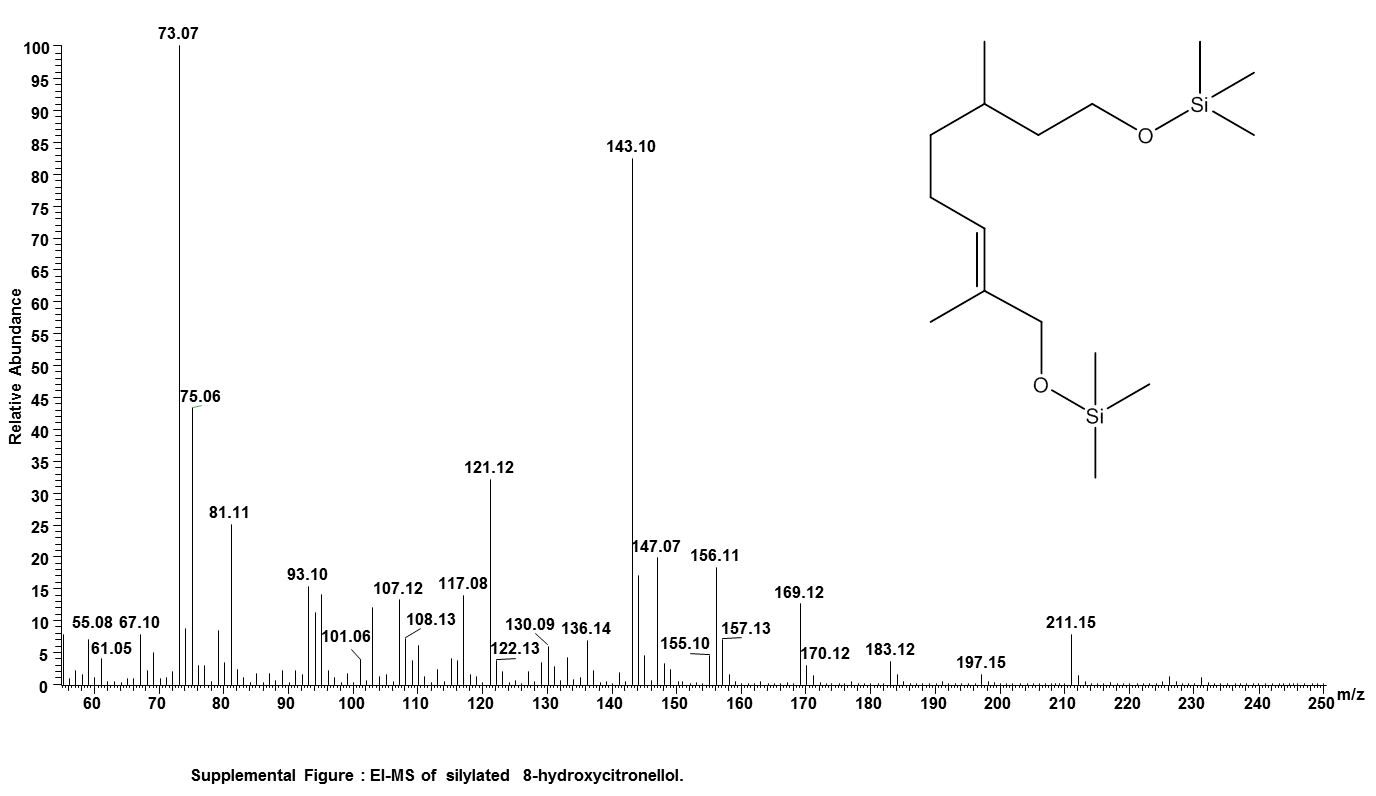

Fig S9 EI-MS of MSTFA silylated 8-hydroxycitronellol. EI mass spectrum corresponding to the peak of 8-hydroxycitronellol formed by CYP6BH5 in Figure 6C. According to NMR data acquired from the non-silylated compound, the peak is identical with 8-hydroxycitronellol.

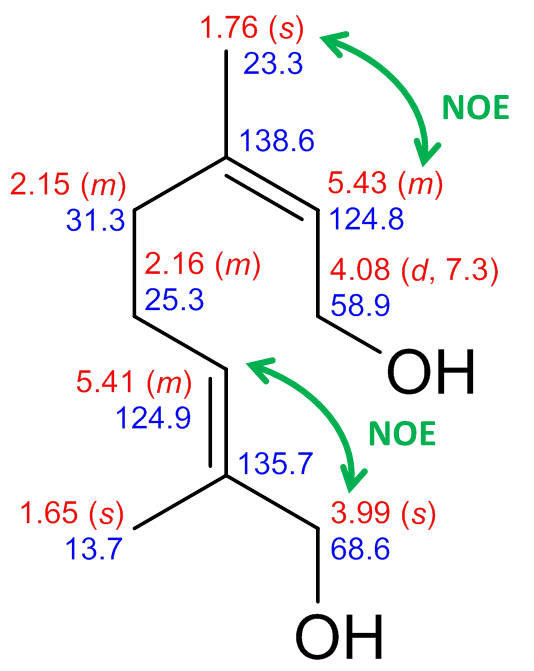

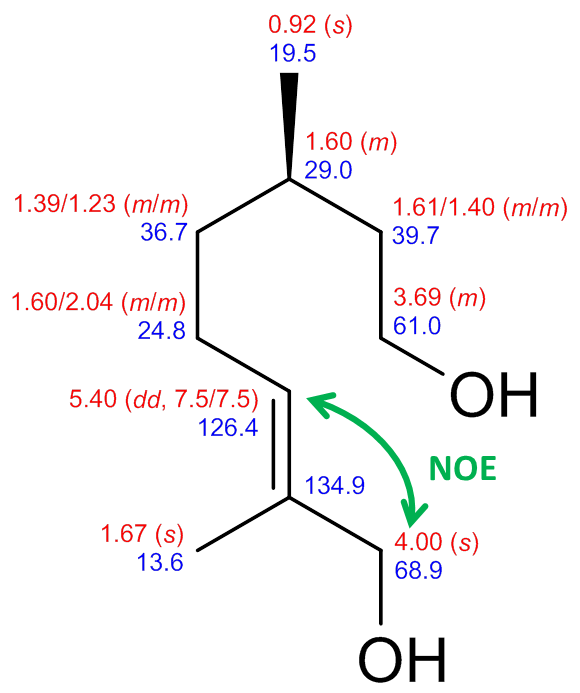

b. 8-OH-citronellol

a. 8-OH-nerol

Fig S10: Structures of compounds formed from nerol and citronellol, respectively, by CYP6BH5. Chemical shifts with multiplicities (δ in ppm, J in Hz, red: ^1^H, blue: ^13^C) were given next to the formulae. Characteristic NOE correlations are indicated in green to determine the stereochemistry.

Data deposition: The CYP6BH5 and CpCPR sequence reported in this paper have been deposited in the GenBank database with accession no. MK843790 and accession no. MK843791 respectively.
